# cGAS–STING induced IFN-β acts as a dual regulator of osteoclastogenesis via direct and osteoblast-mediated mechanisms

**DOI:** 10.64898/2026.05.09.724040

**Authors:** Hannah Simonis, Sanya Middha, Loris Graf, Ramin Naibi, Vivian Polenz, Katharina F. Kubatzky, Elisabeth Seebach

## Abstract

Osteolytic bone diseases are driven by excessive osteoclast formation and bone resorption. While cGAS–STING signaling is known to regulate bone homeostasis via macrophage-intrinsic mechanisms, its role in osteoblast-mediated control of osteoclastogenesis remains poorly defined. Here, we show that cGAS–STING activation of macrophages suppresses their osteoclastogenic potential while promoting immune activation. In osteoblasts, cGAS–STING triggers IRF3-mediated IFN-β production and, notably, shifts the OPG–RANKL axis toward increased osteoprotegerin. In transwell co-culture, pre-activated osteoblasts reduce osteoclast differentiation of strain-matched macrophages. Mechanistically, osteoblast-derived IFN-β is sufficient to inhibit osteoclastogenesis in a paracrine manner. Furthermore, autocrine IFN-β signaling appears to modulate the OPG–RANKL axis, although additional regulatory factors may contribute. Together, these findings identify cGAS–STING-IFN-β signaling as a dual regulator of osteoclastogenesis, acting directly on macrophages and indirectly via osteoblast-derived anti-osteoclastogenic mediators. This highlights osteoblasts as cGAS–STING-responsive bystander cells within the bone microenvironment that can be targeted as an alternative strategy to limit pathological bone resorption.

**GRAPHICAL ABSTRACT:** 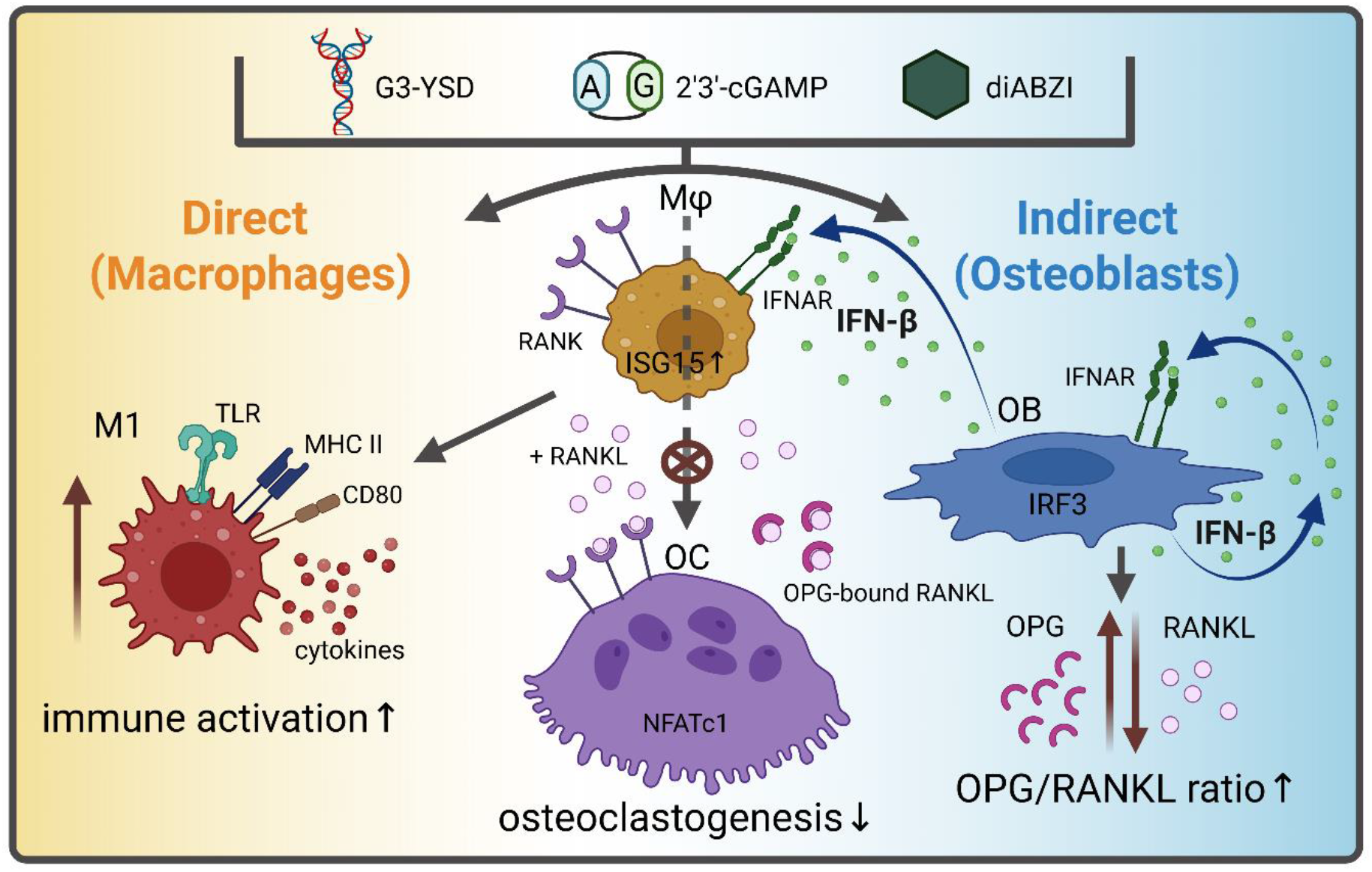

## INTRODUCTION

Bone homeostasis is maintained through the coordinated activity of osteoclasts, which resorb bone, and osteoblasts, which mediate bone formation [1]. Under physiological conditions, the balanced interplay between these cell types ensures continuous bone remodeling and preservation of skeletal integrity. Disruption of this balance, particularly through excessive osteoclast activity, contributes to the development of osteolytic bone diseases such as osteoporosis and inflammation-, infection- and cancer-associated bone loss. These conditions are characterized by progressive bone destruction, elevated fracture risk and a substantial clinical burden. Current therapeutic strategies primarily target osteoclast differentiation and function using RANKL inhibitors (e.g., denosumab) and anti-resorptive agents such as bisphosphonates [2, 3]. While effective in reducing bone resorption, these approaches are associated with impaired bone remodeling and reduced bone quality upon long-term use or discontinuation, highlighting the need for alternative treatment strategies [4-6]. Given the close interplay between the immune system and bone homeostasis, immune modulation has emerged as a promising approach to control pathological bone loss [7, 8]. Targeting pathways that integrate inflammatory signaling with osteoclast differentiation may enable a more refined and physiologically relevant regulation of bone turnover while reducing systemic side effects.

Osteoclasts are multinucleated cells derived from the monocyte–macrophage lineage, and their differentiation is tightly controlled by signals from macrophage colony-stimulating factor (M-CSF) and receptor activator of nuclear factor κB ligand (RANKL), which engages its receptor RANK on osteoclast precursors. Downstream of RANK signaling, the master transcription factor NFATc1 drives osteoclastogenesis by inducing genes required for cell fusion and bone resorption [9, 10]. In contrast, interferon regulatory factor 8 (IRF8) acts as a key negative regulator by suppressing NFATc1 expression and activity, thereby limiting osteoclast differentiation [11]. Type I interferons, in particular IFN-β, further function as anti-osteoclastogenic mediators that restrain osteoclast formation through feedback inhibition of RANK signaling [12, 13]. The balance between these opposing pathways critically determines osteoclastogenic potential.

Given that IFN-β represents a key regulator of osteoclastogenesis, innate immune pathways that control interferon production have emerged as potential modulators of bone homeostasis [14]. The cyclic GMP–AMP synthase (cGAS)–stimulator of interferon genes (STING) pathway is a central component of innate immune signaling that detects cytosolic DNA of self and foreign origin and generates the second messenger 2′3′-cyclic GMP–AMP (2′3′-cGAMP). This second messenger activates STING and induces IRF3-dependent production of IFN-β [15, 16]. Secreted IFN-β subsequently signals in an autocrine and paracrine manner via the type I interferon receptor (IFNAR), leading to activation of downstream JAK–STAT signaling and induction of interferon-stimulated genes such as ISG15 [17, 18]. Recent studies have implicated the cGAS–STING pathway in the regulation of bone homeostasis, primarily through macrophage-intrinsic mechanisms that mediate anti-osteoclastogenic effects via interferon induction and its downstream signaling [19-21]. However, in addition to interferon responses, cGAS–STING activation also engages pro-inflammatory pathways, including NF-κB, which can promote macrophage immune activation and thereby enhance bone degradation [22]. This dual role points to a potential trade-off when directly targeting cGAS–STING signaling in macrophages, as the anti-osteoclastogenic effects of pathway activation may come at the cost of enhanced inflammatory macrophage activation and its potentially detrimental consequences for bone homeostasis. At the same time, cGAS–STING signaling activates multiple cell types within the bone microenvironment that can indirectly influence osteoclast differentiation, raising the possibility that selectively targeting such bystander cells may provide a more refined strategy to control pathological osteoclastogenesis.

In this context, osteoblasts are of particular interest, as they regulate osteoclast differentiation through the secretion of RANKL and its decoy receptor osteoprotegerin (OPG). The ratio of RANKL to OPG represents a central checkpoint in osteoclastogenesis, with increased OPG limiting osteoclast differentiation and bone resorption [23-25]. A recent study showed that systemic activation of STING, achieved by intraperitoneal administration of the synthetic agonist diABZI, alleviates bone loss in both ovariectomy (OVX)-induced osteoporosis and particle-induced calvarial osteolysis models and is associated with a decreased serum RANKL/OPG ratio [14]. In addition, STING deficiency resulted in reduced OPG and increased RANKL levels in mouse tibiae [20]. However, these observations were not further mechanistically addressed, and a direct link between STING signaling and osteoblast function has not been established. Thus, it remains unclear whether osteoblasts are responsive to cGAS–STING signaling and, if so, whether this directly regulates OPG and RANKL production to influence osteoclastogenesis via osteoblast–osteoclast crosstalk. In particular, the contribution of interferon signaling, especially IFN-β, to mediating these potential effects in osteoblasts has not been defined.

Notably, the cGAS–STING pathway can be modulated using synthetic agonists and inhibitors [26]. Double-stranded DNA mimetics such as G3-YSD activate cGAS, whereas small molecules such as RU.521 inhibit cGAS activity. Downstream, STING can be directly activated by cyclic dinucleotides such as 2′3′-cGAMP or diABZI, while inhibitors such as H-151 block STING signaling. These tools provide the opportunity to study cGAS–STING-dependent effects on osteoclastogenesis, inflammatory activation of macrophages, and osteoblast-mediated regulation of bone homeostasis under defined conditions.

Here, we investigated how modulation of cGAS–STING signaling impacts osteoclastogenesis and macrophage immune activation, and whether this pathway can be used to control osteoclast formation indirectly via osteoblasts. Using murine macrophage models, including RAW 264.7 cells and primary bone marrow–derived macrophages (BMDMs), we assessed the effects of cGAS–STING agonists and inhibitors on osteoclast differentiation and inflammatory activation. To address osteoblast–osteoclast crosstalk, we established a transwell co-culture system using primary osteoblasts isolated from long bones and strain-matched BMDMs to determine whether activation of cGAS–STING signaling in osteoblasts modulates osteoclastogenesis in a paracrine manner and alters the OPG–RANKL axis. Furthermore, we investigated the role of osteoblast-derived IFN-β in mediating these effects.

## RESULTS

### Modulation of cGAS–STING signaling in macrophages affects osteoclastogenesis, with activation reducing and early inhibition enhancing osteoclast formation in a cell source-dependent manner

To assess the direct impact of cGAS–STING pathway modulation on osteoclast formation, macrophages were treated with agonists or inhibitors in the presence of the osteoclast-inducing cytokine RANKL. Both primary BMDMs and the well-established macrophage cell line RAW 264.7 were included. We first confirmed expression of cGAS and STING in both cell types. While both proteins were detected in BMDMs and RAW 264.7 cells, STING levels were higher in BMDMs. In contrast, RAW 264.7 cells exhibited multiple bands for cGAS protein, with a more intense full-length band compared to BMDMs, and significantly higher cGAS mRNA copy numbers (Suppl. Figure S1). Activation of cGAS by G3-YSD significantly reduced osteoclast numbers in BMDMs compared to the transfection agent control, whereas no effect was observed in RAW 264.7 cells (Figure 1A). This was associated with IFN-β signaling, as reflected by increased *Isg15* expression in BMDMs upon RANKL stimulation, which was diminished in RAW 264.7 cells (Figure 1B). Consistently, RAW 264.7 cells exhibited reduced gene expression of IRF8 alongside increased gene expression of NFATc1 and downstream osteoclast-associated genes, including *Acp5* (tartrate-resistant acid phosphatase, TRAP), *Atp6v0d2* (d2 isoform of vacuolar ATPase V_0_ domain) and *Ctsk* (Cathepsin K), which were not suppressed and instead tended to be slightly elevated upon cGAS activation (Figure 1C). In additional experiments, pre-activation of cGAS with G3-YSD prior to RANKL stimulation, followed by removal of the ligand before induction of osteoclastogenesis, resulted in a reduction of osteoclast formation by RAW 264.7 cells comparable to that observed in BMDMs (Suppl. Figure S2A). In contrast to co-stimulation conditions, this setup did not lead to a suppression of the cGAS-mediated *Isg15* expression by RANKL and was associated with reduced induction of osteoclast-associated genes (Suppl. Figure S2B+C). In this experimental panel, co-stimulation led to a modest reduction in osteoclast formation also in RAW 264.7 cells. However, control experiments revealed that the YSD control itself affected the macrophage response, therefore, the transfection reagent LyoVec™ alone was used as control in subsequent experiments. Inhibition of cGAS using RU.521 increased osteoclast formation especially in RAW 264.7 cells (Figure 1D). This was accompanied by reduced IFN signaling, as reflected by decreased *Isg15* expression, along with reduced *Irf8* levels and increased expression of *Nfatc1* and osteoclast-associated genes (Figure 1E). Notably, expression of *Atp6v0d2*, a marker associated with osteoclast fusion [27], was markedly increased upon cGAS inhibition. The pro-osteoclastogenic effect of cGAS inhibition occurred predominantly during early stages of differentiation (Figure 1F) and was evident even when inhibition was applied prior to RANKL stimulation, as indicated by increased osteoclast numbers and osteoclastogenic gene expression (Figure 1G).

**Figure 1.**
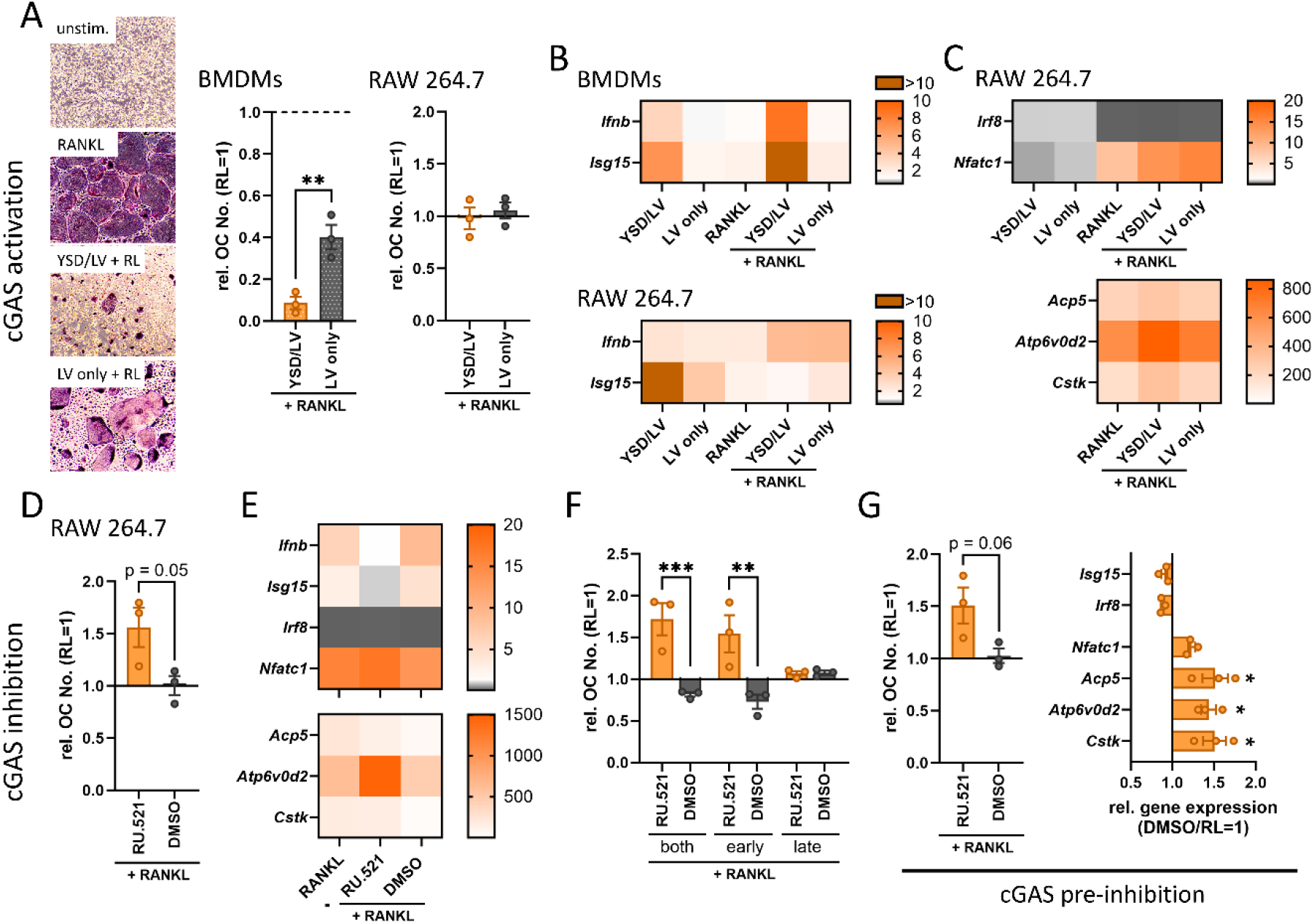
cGAS–STING modulation inversely affects osteoclastogenesis in macrophages. (A) Effect of cGAS activation by G3-YSD complexed in LyoVec™ (YSD/LV; BMDMs: 250 ng/mL, RAW 264.7: 500 ng/mL) on RANKL-mediated osteoclast formation. Representative images of osteoclasts derived from BMDMs (left) and quantification of relative osteoclast numbers per well in BMDMs and RAW 264.7 cells (right). (B+C) Gene expression analysis of interferon-related genes (B) and osteoclast-associated genes (C) 48 h after stimulation with G3-YSD complexed in LyoVec™ (YSD/LV; BMDMs: 250 ng/mL, RAW 264.7: 500 ng/mL) in the presence or absence of 50 ng/mL RANKL. Data are normalized to the unstimulated control. (D–G) Effect of cGAS inhibition using RU.521 (10 µg/mL in DMSO) on osteoclast formation in RAW 264.7 cells. (D) Quantification of relative osteoclast numbers per well. (E) Gene expression analysis of interferon-related and osteoclast-associated genes 48 h after cGAS inhibition in the presence of 50 ng/mL RANKL. Data are normalized to the unstimulated control. (F) Time-dependent effects of cGAS inhibition, with inhibitor (RU.521, 10 µg/mL in DMSO) added throughout differentiation (“both”), during early stages (first 3 days) or during late stages (days 3–5/6). (G) Pre-inhibition of cGAS by treatment with RU.521 (10 µg/mL in DMSO) 24 h prior to RANKL stimulation. The inhibitor was removed before 50 ng/mL RANKL was added. Left: relative osteoclast numbers per well. Right: gene expression analysis of interferon- and macrophage-related genes and osteoclast-associated genes after 24 h cGAS inhibition followed by 48 h RANKL treatment. Data are normalized to the DMSO pre-treated RANKL control. (A-G) BMDMs were cultured in the presence of 25 ng/mL recombinant mouse M-CSF throughout all experiments. Osteoclast numbers per well are shown relatively to the RANKL control. Heatmaps display mean values, and bar graphs show mean ± SEM with individual data points. Statistical analysis was performed using one-way ANOVA with Bonferroni post hoc test (n = 3). RL: RANKL; LV: LyoVec™ transfection agent.

Compared to cGAS, the anti-osteoclastogenic role of STING signaling has already been more extensively investigated, with several studies suggesting that STING pathway activity negatively regulates osteoclast differentiation. In line with these findings, our data show that direct stimulation of STING using the endogenous ligand 2′3′-cGAMP reduced osteoclast formation, with a more pronounced effect in BMDMs and a weaker but still detectable effect when applied at later stages of differentiation in RANKL-primed pre-osteoclasts (Figure 2A). This was accompanied by reduced NFATc1 protein levels upon co-stimulation with RANKL and 2′3′-cGAMP (Figure 2B), increased *Isg15* expression and decreased osteoclast-associated gene expression (Figure 2C). Similarly, the synthetic STING agonist diABZI induced a dose-dependent reduction in osteoclast numbers, again more prominently in BMDMs than in RAW 264.7 cells (Figure 2D). This correlated with a dose-dependent increase in *Isg15* expression, which was further enhanced by RANKL in BMDMs. In contrast, RANKL suppressed STING signaling in RAW 264.7 cells, potentially accounting for the reduced responsiveness in this cell type (Figure 2E). Despite this, STING activation by diABZI also led to a dose-dependent reduction of NFATc1 at both mRNA and protein levels in RAW 264.7 cells, accompanied by decreased expression of osteoclast-associated genes (Figure 2F+G). Inhibition of STING using H-151 resulted in a dose-dependent reduction in osteoclast numbers when applied throughout the entire differentiation period, likely reflecting cytotoxic effects of prolonged H-151 exposure rather than a specific anti-osteoclastogenic mechanism. However, when H-151 was applied only during early stages of osteoclastogenesis, a slight increase in osteoclast formation was observed in BMDMs (Figure 2H).

**Figure 2.**
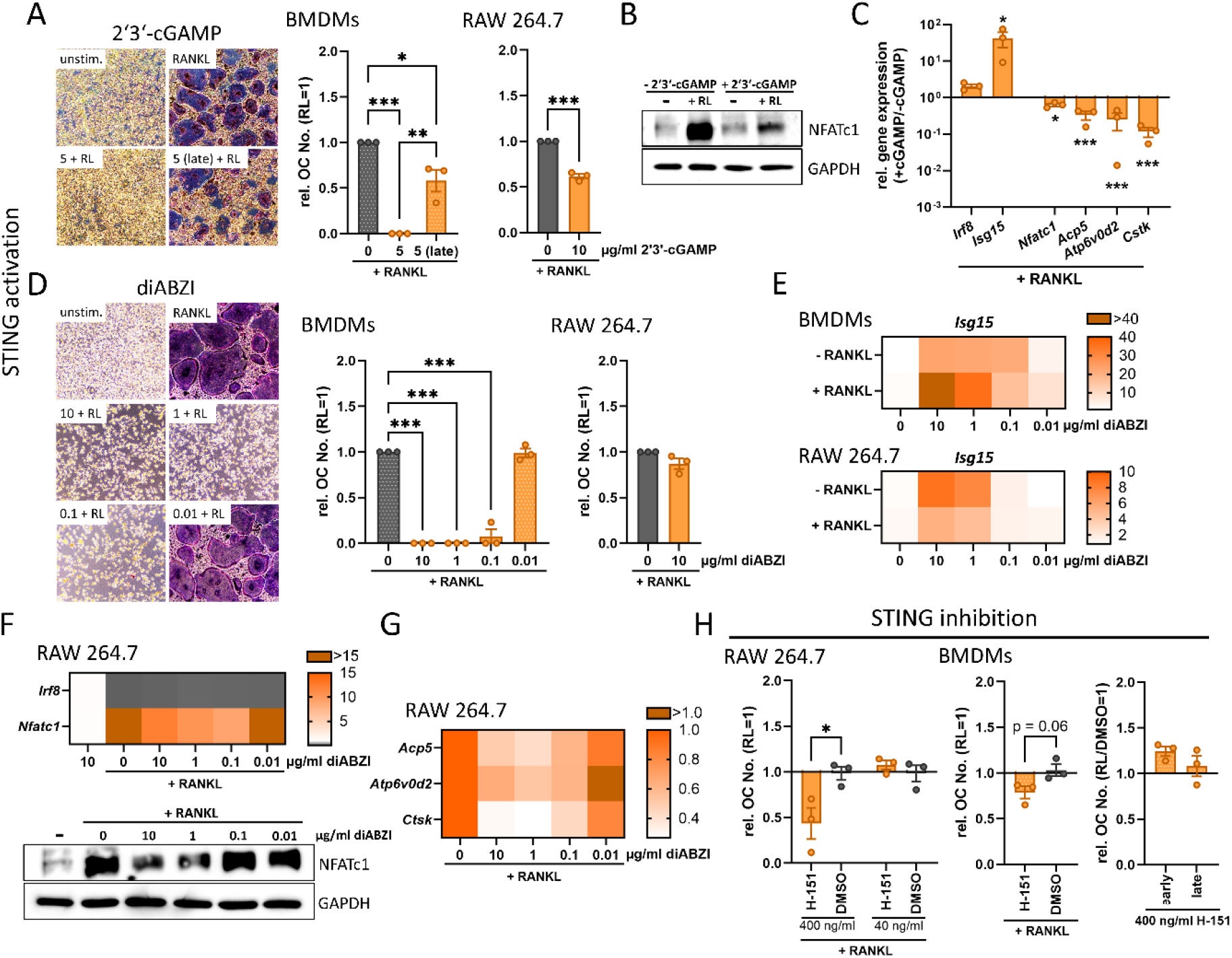
STING modulation inversely affects osteoclastogenesis in macrophages. (A) Effect of STING activation by 2′3′-cGAMP (BMDMs: 5 µg/mL, RAW 264.7: 10 µg/mL) on RANKL-mediated osteoclast formation. 2’3’-cGAMP were given throughout the differentiation or for BMDMs also during late stages (days 3–5/6). Representative images of osteoclasts derived from BMDMs (left) and quantification of relative osteoclast numbers per well in BMDMs and RAW 264.7 cells (right). (B) Immunoblot analysis of NFATc1 protein levels of RAW 264.7 cells following RANKL stimulation (50 ng/mL) in the presence or absence of 10 µg/mL 2′3′-cGAMP. GAPDH served as a loading control. (C) Gene expression analysis of interferon-related, macrophage-related and osteoclast-associated genes of RAW 264.7 cells 48 h after stimulation with 50 ng/mL RANKL with or without 10 µg/mL 2′3′-cGAMP. Data are presented as ratios of +cGAMP to –cGAMP. (D) Effect of STING activation by diABZI (0.01 until 10 µg/mL) on osteoclast formation. Representative images of osteoclasts derived from BMDMs (left) and quantification of relative osteoclast numbers per well in BMDMs and RAW 264.7 cells (right). (E) Induction of the interferon-responsive gene *Isg15* following STING activation with diABZI (0.01 until 10 µg/mL) in BMDMs (upper) and RAW 264.7 cells (lower). Data are normalized to the unstimulated control. (F) Effect of STING activation by diABZI (0.01 until 10 µg/mL) on RANKL-induced NFATc1 expression at mRNA and protein levels after 24 h in RAW 264.7 cells. (G) Gene expression analysis of osteoclast-associated genes 48 h after stimulation with diABZI (0.01 until 10 µg/mL) and 50 ng/mL RANKL in RAW 264.7 cells. Data are normalized to the unstimulated control. (H) Effect of STING inhibition using H-151 (RAW 264.7: 40 or 400 ng/mL in DMSO, BMDMs: 400 ng/mL in DMSO) on osteoclast formation. Left and middle: quantification of relative osteoclast numbers per well upon continuous inhibitor treatment. Right: time-dependent effects of STING inhibition with inhibitor added during early stages (first 3 days) or late stages (days 3–5/6) of differentiation. BMDMs were cultured in the presence of 25 ng/mL recombinant mouse M-CSF throughout all experiments. Osteoclast numbers per well are shown relatively to the RANKL control. Heatmaps display mean values, and bar graphs show mean ± SEM with individual data points. Statistical analysis was performed using one-way ANOVA with Bonferroni post hoc test (n = 3). RL: RANKL.

To further assess the role of IFN-β in this context, BMDMs were stimulated with recombinant IFN-β, which elicited a strong interferon response as reflected by increased *Isg15* expression (Suppl. Figure S3A). Consistently, IFN-β alone was sufficient to suppress osteoclastogenesis at all time points (prior, during early stages or throughout differentiation). Notably, IFN-β treatment also significantly reduced osteoclast numbers when applied at later stages of differentiation, indicating that IFN-β exerts inhibitory effects across multiple phases of osteoclastogenesis (Suppl. Figure S3B). These results are in line with our previous observations in RAW 264.7 cells, where IFN-β similarly suppressed osteoclast formation [28].

Together, these findings demonstrate that cGAS–STING signaling modulates osteoclastogenesis in both primary and immortalized macrophage systems through IFN-β-dependent signaling, while revealing pronounced differences in pathway responsiveness between BMDMs and RAW 264.7 cells. Because cGAS–STING activation is closely coupled to inflammatory signaling, we next investigated to what extent the observed anti-osteoclastogenic effects are accompanied by macrophage immune activation.

### STING activation shifts macrophage fate toward inflammatory activation and away from osteoclastogenic potential

Given that 2′3′-cGAMP elicited the strongest effects in both BMDMs and RAW 264.7 cells and was associated with a marked induction of *Isg15* upon co-stimulation with RANKL, we focused on the impact of STING activation on the macrophage inflammatory response. Stimulation with 2′3′-cGAMP alone increased *Irf8* levels and strongly induced *Ifnb* and *Isg15* expression (Figure 3A). This was accompanied by the release of inflammatory cytokines IFN-β, IL-6, TNF-α and IL-10 (Figure 3B), as well as increased expression of surface activation markers TLR2, MHC class II and CD80 (Figure 3C). This pronounced immune activation was maintained in the presence of RANKL, as indicated by strongly increased MHC class II expression upon combined treatment (Figure 3D). Furthermore, the RANKL-mediated increase in RANK surface expression was attenuated by co-treatment with 2′3′-cGAMP (Figure 3E). Together, these findings indicate that STING activation promotes macrophage immune activation while limiting their osteoclastogenic potential.

**Figure 3.**
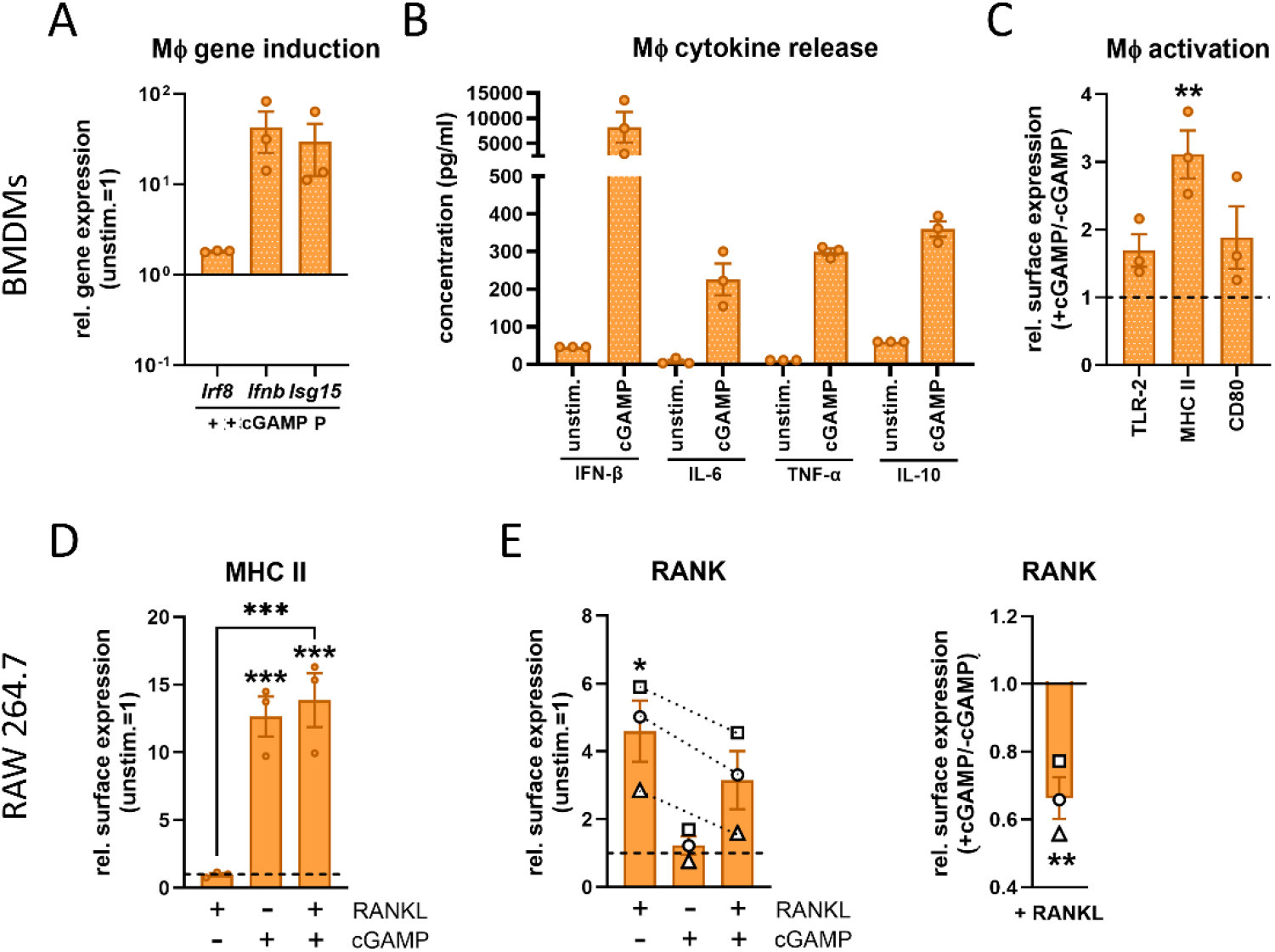
STING activation promotes immune activation while suppressing osteoclastogenic potential in macrophages. (A) Effect of 2′3′-cGAMP stimulation (5 µg/mL) on macrophage- and interferon-related gene expression in BMDMs after 24 h. Data are normalized to the unstimulated control. (B) Cytokine release (IL-6, TNF-α, IL-10 and IFN-β) by 2′3′-cGAMP-stimulated BMDMs after 24 h. (C) Expression of surface activation markers (TLR2, MHC class II and CD80) in BMDMs 24 h after stimulation with 2′3′-cGAMP. Data are presented as ratios of +cGAMP to –cGAMP. (D) MHC class II surface expression in RAW 264.7 cells 24 h after stimulation with 50 ng/mL RANKL, 10 µg/mL 2′3′-cGAMP or the combination of both. Data are normalized to the unstimulated control. (E) RANK surface expression in RAW 264.7 cells 24 h after stimulation with 50 ng/mL RANKL, 10 µg/mL 2′3′-cGAMP or the combination of both. Left: RANK levels normalized to the unstimulated control. Right: RANK expression presented as ratios of +cGAMP to –cGAMP. Different symbols and dotted lines indicating independent experiments, respectively. (A-E) BMDMs were cultured in the presence of 25 ng/mL recombinant mouse M-CSF throughout all experiments. Bar graphs show mean ± SEM with individual data points. Statistical analysis was performed using one-way ANOVA with Bonferroni post hoc test (n = 3). Mϕ: macrophage.

### cGAS–STING activation in osteoblasts modulate the OPG–RANKL axis and drives IFN-β mediated osteoblast–osteoclast crosstalk to suppress osteoclastogenesis

While these findings demonstrate that cGAS–STING activation intrinsically shifts macrophages toward an immune-activated, anti-osteoclastogenic state, cGAS–STING signaling may also affect additional cell types within the bone microenvironment that indirectly regulate osteoclast differentiation. We therefore next investigated whether osteoblasts represent such responsive bystander cells and whether their potential activation through cGAS–STING signaling influences osteoclastogenesis via the release of osteoblast-derived factors. Therefore, we first characterized osteoblasts isolated from long bone fragments of mice. These cells exhibited the expected morphology and were capable of mineralization (Suppl. Figure S4A). They showed increased CD73 expression, a marker of osteoblasts, whereas CD45 expression, indicative of contaminating immune cells, decreased over successive passages, demonstrating enrichment of the osteoblast population, which reached high purity after passage 3 (Suppl. Figure S4B). Furthermore, osteoblasts displayed high STING protein levels, however, cGAS was not visible by immunoblotting, but was clearly detectable on transcript levels (Suppl. Figure S4C+D). Despite this, stimulation with the cGAS agonist G3-YSD revealed that osteoblasts were highly responsive, as indicated by robust induction of *Ifnb, Isg15* and *Il6* expression as well as *cgas* and *Sting1* themselves. Notably, cGAS activation reduced expression of *Tnfsf11*, encoding RANKL, while increasing expression of *Tnfrsf11b*, encoding osteoprotegerin (OPG), resulting in a significant increase in the OPG/RANKL ratio (Figure 4A). Consistent with these findings, elevated levels of IFN-β and OPG were detected in the supernatant following cGAS activation (Figure 4B). In contrast, responses to direct STING activation using 2′3′-cGAMP or diABZI were less pronounced, with reduced changes in gene expression and lower induction of IFN-β (Figure 4A+B). IFN-β levels were moderately increased upon 2′3′-cGAMP stimulation but remained undetectable following diABZI treatment. Instead, OPG levels were highest with 2′3′-cGAMP and comparable between diABZI and G3-YSD (Figure 4B). To determine whether cGAS–STING activation in osteoblasts functionally impacts osteoclastogenesis, we performed transwell co-culture experiments with strain-matched macrophages (BMDMs). Pre-activation of osteoblasts with cGAS–STING agonists for 24 hours significantly reduced osteoclast differentiation of co-cultured macrophages only upon G3-YSD treatment, whereas 2′3′-cGAMP and diABZI showed no effect under these conditions (Figure 4C). To avoid direct effects of residual cGAS–STING agonists on macrophages, osteoblast medium was exchanged prior to co-culture. Given the rapid kinetics of STING activation compared to cGAS-mediated signaling, we reasoned that this procedure may have removed transiently produced soluble factors following direct STING activation. In contrast, the delayed response induced by cGAS activation may allow sustained production of such factors even after medium exchange, thereby preserving their functional impact on osteoclastogenesis. Indeed, STING pathway activation was evident by phosphorylation of IRF3 as early as 3 hours after stimulation with 2’3’-cGAMP or diABZI, but not G3-YSD (Figure 4D), followed by induction of downstream target genes after 6 hours of 2′3′-cGAMP or diABZI treatment (Figure 4E). RUNX2 protein levels remained consistently high across all conditions, indicating preservation of osteoblast identity and differentiation status upon cGAS–STING activation. At this time point, *Tnfrsf11b* expression was already increased, resulting in an elevated OPG/RANKL ratio, although no reduction in *Tnfsf11* expression was yet observed. Consistent with this kinetics, shortening the pre-activation period restored the functional effect of direct STING activation in co-culture, with both 2′3′-cGAMP and diABZI significantly reducing osteoclast formation (Figure 4F). The effect was more pronounced for 2′3′-cGAMP, suggesting differences in uptake kinetics (diffusion versus transporter-mediated) or downstream signaling between the agonists. Together, these findings indicate that osteoblast-derived factors released upon cGAS–STING activation suppress macrophage osteoclastogenesis in a paracrine manner and may also modulate the OPG–RANKL axis through autocrine signaling. Based on the robust induction of IFN-β release from osteoblasts, we next asked whether IFN-β signaling mediates these anti-osteoclastogenic effects.

**Figure 4.**
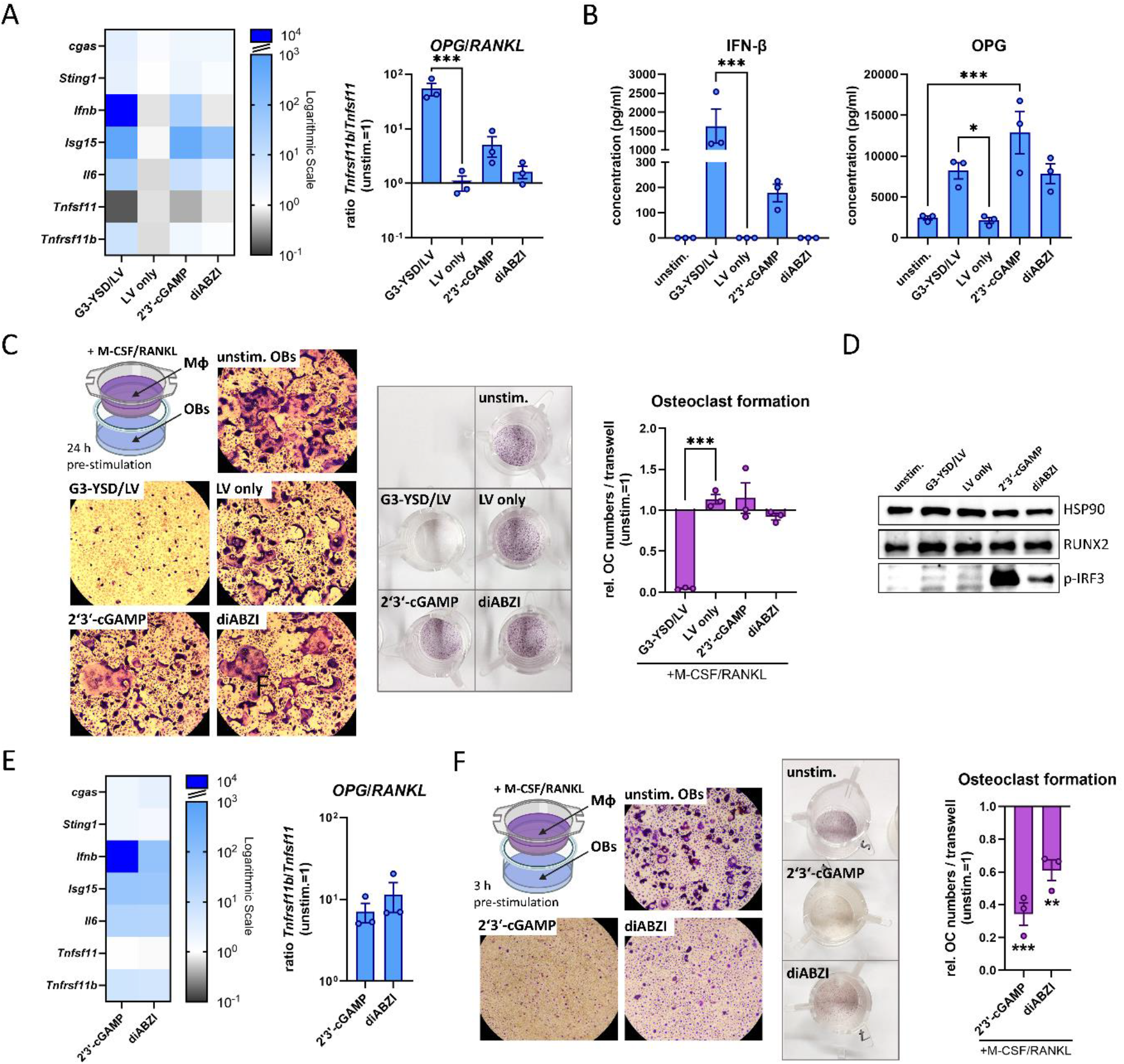
cGAS–STING activation in osteoblasts induces IFN-β production, modulates the OPG–RANKL axis and suppresses osteoclast differentiation of macrophages in co-culture. (A) Gene expression analysis of osteoblasts after stimulation with cGAS–STING agonists for 24 h (G3-YSD: 500 ng/mL, 2’3’-cGAMP: 5 µg/mL, diABZI: 500 ng/mL). Left: heatmap of cGAS–STING pathway-associated genes, including *Tnfsf11* (RANKL) and *Tnfrsf11b* (OPG). Right: ratio of *Tnfrsf11b* to *Tnfsf11* mRNA levels. Data are normalized to the unstimulated control. (B) Protein concentrations of IFN-β and OPG in supernatants of osteoblasts after 24 h stimulation with cGAS–STING agonists (G3-YSD: 500 ng/mL, 2’3’-cGAMP: 5 µg/mL, diABZI: 500 ng/mL). (C) Co-culture model with osteoblasts seeded in the lower compartment and pre-stimulated with cGAS–STING agonists for 24 h (G3-YSD: 500 ng/mL, 2’3’-cGAMP: 5 µg/mL, diABZI: 500 ng/mL), followed by co-culture with strain-matched BMDMs in transwell inserts and induction of osteoclastogenesis using 25 ng/mL M-CSF and 50 ng/mL RANKL. Osteoclast formation was assessed by TRAP staining. Left: representative images of osteoclasts. Middle: images of transwell inserts after TRAP staining. Right: quantification of osteoclast numbers per transwell. (D) Immunoblot analysis of RUNX2 protein levels and IRF3 pathway activation, indicated by phosphorylated IRF3, in osteoblasts after 3 h of cGAS– STING stimulation (G3-YSD: 500 ng/mL, 2’3’-cGAMP: 5 µg/mL, diABZI: 500 ng/mL). HSP90 served as a loading control. (E) Gene expression analysis of osteoblasts after stimulation with STING agonists for 6 h (2’3’-cGAMP: 5 µg/mL, diABZI: 500 ng/mL). Heatmap and OPG–RANKL ratio are presented as described in (B). (F) Co-culture model as described in (C), with osteoblasts pre-stimulated with STING agonists (2’3’-cGAMP: 5 µg/mL, diABZI: 500 ng/mL) for 3 h prior to co-culture with BMDMs. (A-F) Osteoclast numbers per well are shown relatively to the unstimulated osteoblast control. Heatmaps display mean values, and bar graphs show mean ± SEM with individual data points. Statistical analysis was performed using one-way ANOVA with Bonferroni post hoc test (n = 3). OB: osteoblast; OC: osteoclast; Mϕ: macrophage; LV: LyoVec™ transfection agent.

To test whether osteoblast-derived IFN-β contributes to autocrine modulation of osteoblast function, we pre-stimulated osteoblasts with recombinant IFN-β. This resulted in a slight reduction in osteoclast numbers in co-culture (Figure 5A), which was less pronounced than the effects observed following cGAS–STING activation with G3-YSD or 2′3′-cGAMP but comparable to diABZI treatment. Osteoblasts responded to IFN-β stimulation with increased expression of *Isg15* and *Il6*, and notably, *Tnfrsf11b* mRNA levels were elevated, whereas *Tnfsf11* expression was only modestly reduced. Overall, this resulted in an increased OPG/RANKL ratio upon IFN-β treatment (Figure 5B). To further assess the role of IFN-β in osteoblast responses downstream of STING activation, we neutralized IFNAR1 and stimulated osteoblasts with 2′3′-cGAMP or diABZI. IFN signaling was not completely abolished but was clearly reduced, as indicated by decreased *Isg15* expression in the IFNAR1-blocked group compared to IgG controls (Suppl. Figure S5, left). Notably, the increase in the OPG/RANKL ratio upon STING activation was also attenuated (Suppl. Figure S5, right). This effect became more apparent when comparing IFNAR1 blockade to control antibody conditions, revealing reduced expression of IFN-responsive genes, increased *Tnfsf11* expression, and decreased *Tnfrsf11b* mRNA levels, ultimately resulting in a significantly reduced OPG/RANKL ratio (Figure 5C). These findings indicate that modulation of the OPG–RANKL axis upon STING activation is at least partially dependent on autocrine IFN-β signaling. To determine whether IFN-β is sufficient to mediate the anti-osteoclastogenic effects of osteoblasts, we mimicked the co-culture setup by adding recombinant IFN-β to the lower compartment, while macrophages were cultured with M-CSF/RANKL in transwell inserts. This configuration ensured that IFN-β diffused from the lower chamber to the macrophages, thereby recapitulating paracrine signaling in the absence of osteoblasts. IFN-β significantly reduced osteoclast formation, with an effect size intermediate between cGAS–STING activation by G3-YSD and 2′3′-cGAMP (Figure 5D). Together, these findings identify IFN-β as a dual mediator of osteoblast–osteoclast communication, acting in a paracrine manner to suppress osteoclastogenesis in macrophages while also exerting autocrine effects on osteoblasts to modulate the OPG–RANKL axis, thereby serving as a central driver of the anti-osteoclastogenic response downstream of cGAS–STING activation in osteoblasts.

**Figure 5.**
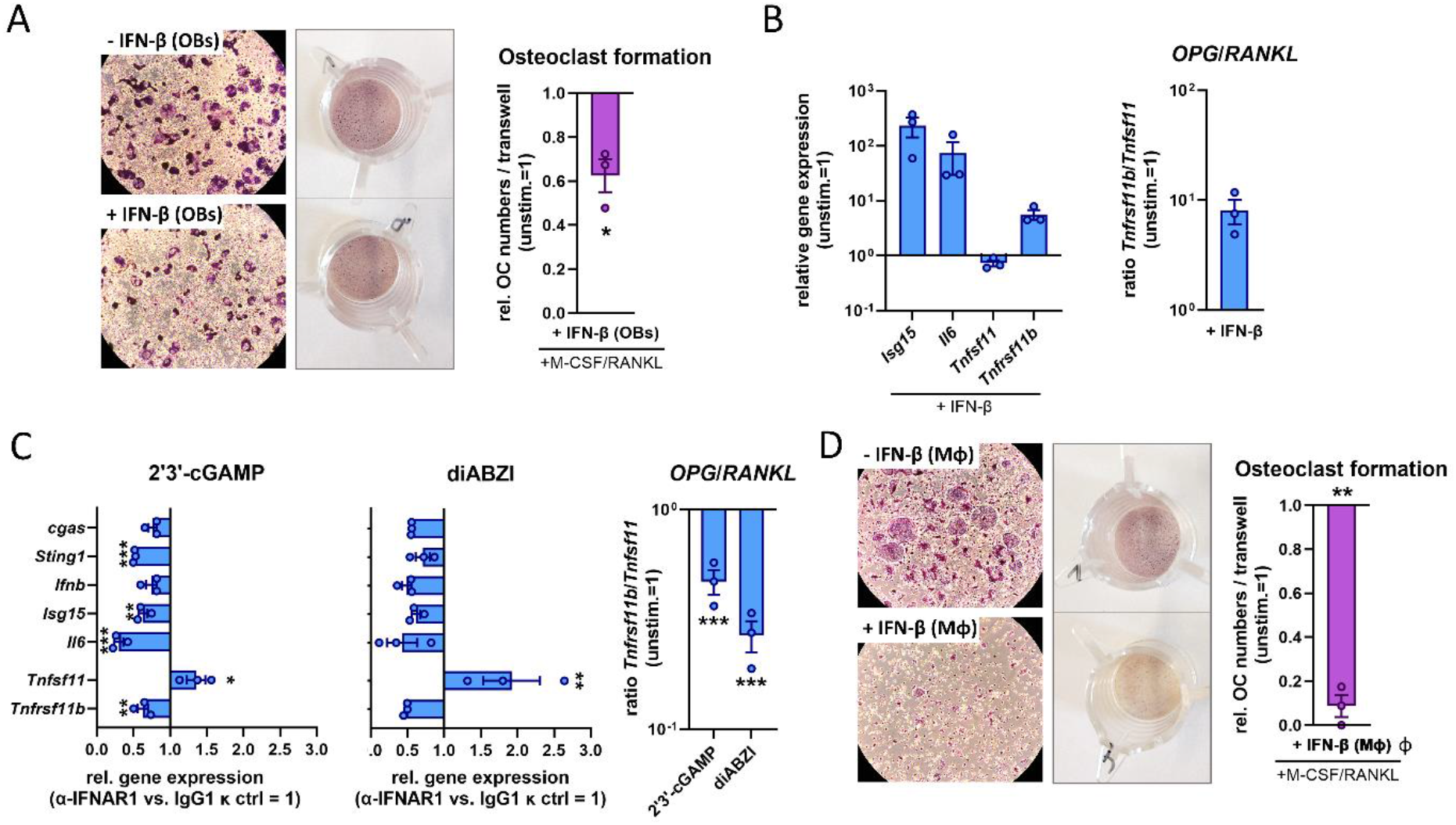
Osteoblast-derived IFN-β modulates the OPG–RANKL axis and is sufficient to inhibit osteoclastogenesis. (A) Co-culture model with osteoblasts seeded in the lower compartment and pre-stimulated with recombinant mouse IFN-β (10 ng/mL) for 3 h, followed by co-culture with strain-matched BMDMs in transwell inserts and induction of osteoclastogenesis using 25 ng/mL M-CSF and 50 ng/mL RANKL. Osteoclast formation was assessed by TRAP staining. Left: representative images of osteoclasts. Middle: images of transwell inserts after TRAP staining. Right: quantification of osteoclast numbers per transwell. (B) Gene expression analysis of osteoblasts after stimulation with recombinant mouse IFN-β (10 ng/mL) for 3 h. Left: expression of IFN-responsive genes, including *Tnfsf11* (RANKL) and *Tnfrsf11b* (OPG). Right: ratio of *Tnfrsf11b* to *Tnfsf11* mRNA levels. Data are normalized to the unstimulated control. (C) Gene expression analysis of osteoblasts after neutralization of IFNAR1 (10 µg/mL) and stimulation with STING agonists (2’3’-cGAMP: 5 µg/mL, diABZI: 500 ng/mL) for 6 h. Relative gene expression is shown as ratios of IFNAR1-neutralized to control antibody (IgG1 κ isotype, both 10 µg/mL) conditions for 2′3′-cGAMP (left) and diABZI (middle). Right: ratio of *Tnfrsf11b* to *Tnfsf11* mRNA levels for both STING agonists. (D) Co-culture model as described in (A), with recombinant mouse IFN-β (10 ng/mL) added to the lower compartment in the absence of osteoblasts. Left: representative images of osteoclasts. Middle: images of transwell inserts after TRAP staining. Right: quantification of osteoclast numbers per transwell. (A-D) BMDMs were cultured in the presence of 25 ng/mL recombinant mouse M-CSF throughout all experiments. Heatmaps display mean values, and bar graphs show mean ± SEM with individual data points. Statistical analysis was performed using one-way ANOVA with Bonferroni post hoc test (n = 3). OB: osteoblast; OC: osteoclast; Mϕ: macrophage; Ctrl: control.

## DISCUSSION AND CONCLUSION

Osteolytic bone diseases are characterized by dysregulated bone homeostasis, in which excessive osteoclast activity drives pathological bone resorption. Although anti-resorptive therapies are available, long-term and systemic suppression of osteoclast function is associated with impaired bone remodeling. This highlights the need for more targeted approaches that do not simply block global bone resorption but instead enable functionally selective and locally controlled regulation of bone turnover. Given the close interplay between the immune system and skeletal homeostasis, immunoregulatory pathways have emerged as promising modulators of osteoclastogenesis. Among these, type I interferons, particularly IFN-β, act as key negative regulators by counteracting RANK signaling in osteoclast precursors. The cGAS–STING pathway is a central upstream regulator of IFN-β production and has been implicated in bone homeostasis, primarily through macrophage-intrinsic mechanisms that suppress their osteoclastogenic differentiation potential. However, because cGAS– STING signaling simultaneously activates pro-inflammatory pathways, direct targeting in macrophages may involve a trade-off between anti-osteoclastogenic effects and enhanced inflammatory activation. In addition, cGAS–STING activation occurs across multiple cell types within the bone microenvironment, raising the possibility that non-osteoclastic cells may indirectly contribute to the regulation of osteoclastogenesis. Here, we investigated both the direct effects of cGAS–STING modulation in macrophages on their osteoclastogenic potential and immune activation, as well as whether osteoblasts represent responsive bystander cells whose activation through cGAS–STING signaling influences osteoclastogenesis via the release of osteoblast-derived anti-osteoclastogenic factors.

Activation of the cGAS–STING pathway in macrophages at both the cGAS and STING level induced a robust type I interferon response, reflected by strong upregulation of the interferon-stimulated gene *Isg15*. Notably, the dynamics of *Isg15* expression differed between the two cell systems. In primary BMDMs, elevated *Isg15* levels were maintained even in the presence of RANKL, whereas in RAW 264.7 cells, concomitant RANKL stimulation attenuated *Isg15* induction unless agonists were applied prior to RANKL exposure. Functionally, *Isg15* expression inversely correlated with osteoclast formation and expression of osteoclastogenic genes, particularly the fusion mediator *Atp6v0d2*, consistent with previous reports identifying ISG15 as a negative regulator of osteoclastogenesis through suppression of ATP6V0D2-dependent cell fusion [29]. Together with recent findings on tonic STING signaling in bone homeostasis [19], our data support a model in which cGAS–STING-dependent interferon signaling represents a central mechanism limiting osteoclast differentiation and maintaining the intrinsic anti-osteoclastogenic state of macrophages. Consistent with this concept, STING agonists were shown to suppress RANKL-induced osteoclastogenesis in macrophages via type I interferon signaling [14, 30]. Our findings align with this, showing that STING activation reduces osteoclast formation. We further demonstrate that upstream cGAS activation and its product, 2′3′-cGAMP, are sufficient to elicit anti-osteoclastogenic effects. However, this anti-osteoclastogenic activity is accompanied by a strong pro-inflammatory response, indicating a prioritization of inflammatory programs over osteoclastogenic commitment. Similar to TLR signaling, the effect depends on the differentiation state: macrophages are more susceptible when activation coincides with early RANKL stimulation, whereas pre-committed osteoclast precursors are less affected [28, 31, 32]. In line with this, 2′3′-cGAMP was less effective at later stages of RANKL-induced differentiation, consistent with a report on bacterial cyclic dinucleotides in murine BMMs [30]. The role of STING inhibition in osteoclastogenesis is less consistent. While STING deficiency had been linked to increased osteoclast formation and reduced bone mass [19, 20], pharmacological inhibition could reduce differentiation by suppressing NF-κB and inflammatory pathways [33]. These effects were temporally dependent: early inhibition reduced osteoclast formation, whereas later inhibition enhanced differentiation, reflecting dynamic type I interferon and *Isg15* responses [34]. Timing also affected STING activation, with opposing effects at different stages. These findings differ from our observations, as well as from previous reports demonstrating predominantly anti-osteoclastogenic effects of STING activation. Overall, these data show that direct modulation of cGAS–STING in macrophages produces complex, context-dependent effects, with anti-osteoclastogenic outcomes coupled to pro-inflammatory activation and strongly influenced by differentiation state of the cells. This suggests that targeting macrophages directly may have limited therapeutic potential, highlighting the value of alternative, cGAS–STING-responsive bystander cells. In this context, osteoblasts are an attractive candidate, as key regulators of osteoclastogenesis via the OPG–RANKL axis and mediators of indirect, paracrine effects.

Most studies have focused on cGAS–STING activation in macrophages, with comparatively little attention on osteoblasts. Previous work reported that STING deficiency impairs osteoblastogenesis, reducing RUNX2 levels in tibiae and ALP activity in bone marrow stromal cells [20], whereas direct STING activation had little measurable effect in a co-culture system of BMMs and calvarial mouse osteoblasts, likely due to low or undetectable STING expression in the osteoblasts used [30]. This is consistent with preliminary data of our group, where STING was undetectable in the calvarial osteoblast cell line MC3T3 and STING activation did not affect osteoclastogenesis in our model (data not shown). In contrast, primary mouse osteoblasts isolated from long bones expressed robust STING, whereas cGAS levels were low or barely detectable. Despite minimal cGAS expression, these cells responded to the cGAS agonist G3-YSD and the STING agonists 2′3′-cGAMP and diABZI with a strong IRF3-mediated type I interferon response, with differences in activation kinetics likely reflecting distinct uptake mechanisms—G3-YSD requires transfection, 2′3′-cGAMP relies on transporters, and diABZI enters via passive diffusion. The subsequent release of IFN-β by osteoblasts directly inhibited osteoclast formation in co-culture with macrophages, demonstrating a paracrine anti-osteoclastogenic effect. Importantly, and to our knowledge for the first time, we demonstrate that cGAS–STING activation in osteoblasts also led to a downregulation of *Tnfsf11* (RANKL) and a concomitant upregulation of *Tnfrsf11b* (OPG). This shift in the OPG–RANKL balance was further supported by significantly increased OPG secretion, highlighting a potent osteoblast-mediated suppression of osteoclastogenesis upon cGAS-STING signaling. Previous studies had reported STING-dependent changes in OPG and RANKL—STING deficiency was associated with reduced OPG and increased RANKL expression in mouse tibiae [20], while STING activation decreased the RANKL/OPG ratio in the serum of OVX-induced osteoporotic mice [14]—but the cellular source and underlying molecular mechanisms were unclear. Furthermore, our data identify IFN-β as a key mediator connecting cGAS–STING activation to OPG induction in osteoblasts. Beyond its direct inhibition of osteoclast differentiation, IFN-β drives OPG expression, thereby indirectly modulating osteoclastogenesis via the RANKL–OPG axis. Consistent with this, clinical observations in multiple sclerosis patients treated with IFN-β showed time-dependent changes in circulating RANKL and OPG [35]. Collectively, these findings reveal osteoblasts as a previously underrecognized target of cGAS–STING activation, with osteoblast-derived IFN-β acting as a central mediator that both inhibits osteoclast differentiation in a paracrine manner and autocrinely induces OPG expression in osteoblasts, thereby shifting the OPG–RANKL axis towards an anti-osteoclastogenic state.

A limitation of our study is that the specific contribution of OPG to the observed effects of cGAS-STING pre-activated osteoblasts on RANKL-mediated osteoclastogenesis was not directly addressed. Recent studies demonstrate that OPG acts predominantly in a local manner and that osteoblasts represent the functional source of OPG in bone [36, 37]. Notably, osteoblast-derived OPG is essential for maintaining bone homeostasis and directly suppresses osteoclastogenesis by limiting RANKL activity, underscoring the importance of local RANKL–OPG regulation within the bone microenvironment. These findings suggest that osteoblast-derived OPG may substantially contribute to the anti-osteoclastogenic effects observed in our system. Given our data linking IFN-β signaling to the regulation of OPG expression in osteoblasts, it is conceivable that OPG functions as a key downstream effector that complements the inhibitory effects of IFN-β on osteoclasts. Furthermore, the observation that OPG induces STING expression in macrophages upon RANKL stimulation [38] raises the possibility of a synergistic activation loop involving STING-dependent IFN signaling, through which OPG and IFN-driven pathways may cooperatively enhance the inhibition of osteoclast formation. This putative crosstalk and its contribution to bone homeostasis will require further investigation.

Taken together, our findings extend the current understanding of cGAS–STING signaling in bone biology by revealing a cell type–dependent dichotomy in its functional output: direct activation in macrophages suppresses osteoclast differentiation—consistent with the inhibitory role of IFN-β on RANK signaling—but is accompanied by a pronounced pro-inflammatory response that may limit its therapeutic potential in chronic bone diseases. In contrast, osteoblasts act as cGAS–STING-responsive bystander cells, in which pathway activation elicits a robust IFN-β response without requiring direct immune cell activation. Osteoblast-derived IFN-β inhibits osteoclast formation in a paracrine manner and, in an autocrine manner, reinforces osteoblast function by shifting the OPG–RANKL balance toward an anti-osteoclastogenic profile. By engaging these bystander cells, cGAS–STING signaling can indirectly regulate osteoclastogenesis while avoiding the pro-inflammatory consequences of direct macrophage activation. Targeting this pathway in osteoblasts may therefore provide a more selective and therapeutically favorable strategy, combining the anti-osteoclastogenic activity of IFN-β with modulation of the OPG–RANKL axis to achieve a refined control of bone turnover.

## RESOURCE AVAILABILITY

### Lead contact

Further information and requests for resources and reagents should be directed to and will be fulfilled by the lead contact, Dr. Elisabeth Seebach.

### Materials availability

This study did not generate new unique reagents.

### Data and code availability

All data supporting the findings of this study are available from the lead contact upon reasonable request.

## ACKNOWLEDGEMENT

We would like to thank Magdalena Urlaub for help with the cell culture and experiments as well as Gabriele Sonnenmoser and Jannis Pliatsikos for technical support. This research did not receive any specific grant from funding agencies in the public, commercial, or not-for-profit sectors. The graphical abstract was created in BioRender (Seebach, E. (2026) https://BioRender.com/eu8dyeq).

## AUTHOR CONTRIBUTIONS

HS, SM, LG, RN and VP participated in data acquisition and analysis. KFK contributed to data interpretation and critically revised the manuscript. ES was responsible for study conception, interpretation of data and wrote the manuscript. All authors read and approved the final manuscript.

## DECLARATION OF GENERATIVE AI AND AI-ASSISTED TECHONOLOGIES IN THE WRITING PROCESS

During the preparation of this manuscript, ChatGPT was used for English language editing and Elicit was used to assist with literature discovery. After using this tool/service, the authors reviewed and edited the content as needed and take full responsibility for the content of the published article.

## METHODS

## EXPERIMENTAL MODEL AND SUBJECT DETAILS

### Mice

Female adult C57BL/6N wild-type mice (age: 3 to 6 month) were used for isolation of primary cells. Animals were purchased from Janvier Labs (Le Genest-Saint-Isle, France) and maintained under specific pathogen–free (SPF) conditions. All animal procedures were conducted in accordance with national guidelines for animal care and were approved by the local authorities (T-38/24).

### Primary cells

#### Bone marrow–derived macrophages (BMDMs)

Bone marrow cells were isolated from femurs and tibiae by centrifugation (5000 rpm for 2 min) as previously described (REF) and cultured in high-glucose DMEM supplemented with 10% heat-inactivated FCS, 1% penicillin/streptomycin, 50 μM β-mercaptoethanol (BMDM medium), and L929 cell-conditioned medium (LCCM). Fresh LCCM was added on day 4, and differentiated macrophages were used on day 7. In experiments, BMDM medium was supplemented with 25 ng/mL recombinant mouse M-CSF (Bio-Techne Ltd., USA).

#### Long bone–derived osteoblasts (LBOBs)

Primary osteoblasts were isolated from femurs and tibiae following removal of bone marrow. Bones were cut into small fragments and digested with 2 mg/mL collagenase II (PAN-Biotech) in culture medium for 2 h at 37°C. Bone fragments were washed twice and transferred to tissue culture–treated dishes containing α-MEM supplemented with 10% heat-inactivated fetal calf serum (FCS), 1% penicillin/streptomycin, 2 mM GlutaMAX™ (Thermo Fisher Scientific) and 10 ng/mL ascorbic acid (Sigma-Aldrich) (OB medium). Osteoblasts were allowed to grow out from bone fragments, which were subsequently removed. Culture medium was replaced every 3 days. Cells were expanded to ∼80% confluency and passaged by trypsinization. Osteoblast identity and purity were verified by assessing absence of hematopoietic contamination (CD45) and enrichment of osteoblastic cells (CD73) and by evaluation of mineralization capacity using Alizarin Red staining. Cells were used for experiments between passages 4 and 6.

#### Cell lines

The murine macrophage cell line RAW 264.7 (ATCC TIB-71, USA) was cultured in high-glucose DMEM supplemented with 10% heat-inactivated fetal calf serum (FCS) and 1% penicillin/streptomycin at 37°C and 5% CO2. Cells were passaged every 2-3 days by scraping. Only day 2 cultures between passage 5-25 and without any signs of pre-activation were used for experiments.

## METHOD DETAILS

### Macrophage stimulation and osteoclast differentiation

RAW 264.7 cells and BMDMs were seeded at densities depending on the downstream application. For osteoclast differentiation assays, RAW 264.7 cells were seeded at 5 × 10^3^ cells/well in 24-well plates and BMDMs at 7 × 10^4^ cells/well in 48-well plates (1 mL medium per well). For gene expression analyses, RAW 264.7 cells were seeded at 1 × 10^5^ cells/well in 24-well plates and BMDMs at 1 × 106 cells/well in 12-well plates (1 mL medium per well). For immunoblotting, RAW 264.7 cells (0.5–1 × 106 cells/well) and BMDMs (2 × 106 cells/well) were seeded in 6-well plates (2 mL medium per well).

Cells were stimulated with 50 ng/mL recombinant mouse RANKL (Bio-Techne Ltd., USA) to induce osteoclast differentiation. BMDMs were additionally supplemented with 25 ng/mL M-CSF. Where indicated, cells were treated with: cGAS agonist G3-YSD (RAW 264.7: 500 ng/mL; BMDMs: 250 ng/mL) complexed with LyoVec™ (1:100, 15 min pre-incubation), cGAS inhibitor RU.521 (10 µg/mL in DMSO) added 3 h prior to stimulation; STING agonists 2′3′-cGAMP (RAW: 10 μg/mL; BMDMs: 5 μg/mL) or diABZI (0.01–10 μg/mL), STING inhibitor H-151 (40 or 400 ng/mL in DMSO) added 2 h prior to stimulation (all InvivoGen, USA). Vehicle controls (DMSO, LyoVec™) were included in all experiments. For molecular analyses, cells were harvested at indicated time points (3 or 24 h for protein analysis; 6, 24, 48 or 72 h for gene expression). For osteoclast formation, cells were re-stimulated on day 3 by replacing half of the medium and adding fresh stimuli at half of the initial concentration. Osteoclast numbers were assessed on days 5–6. In experiments with RAW 264.7 cells extending beyond 3 days, the medium was supplemented with 50 µM β-mercaptoethanol to limit proliferation and prevent overgrowth of the cell line. To distinguish effects on early versus late osteoclastogenesis, cGAS-STING agonists or inhibitors were applied either during the first 3 days or from day 3 to days 5–6. For these experiments, the medium was completely replaced at day 3, and M-CSF and RANKL were added at full concentration. For cGAS pre-inhibition, RAW 264.7 cells were treated with the inhibitor RU.521 (10 µg/mL in DMSO) or DMSO alone for 24 h. After medium exchange, RANKL was added to induce osteoclastogenesis.

### Osteoblast stimulation

Before experiments, osteoblasts were seeded in the respective well format overnight (1 × 10^5^ cells/well in 12-well plate for gene expression; 2 × 10^5^ cells/well in 6-well plate for protein analysis). The next day, medium was exchanged and osteoblasts were stimulated with cGAS–STING agonists (2′3′-cGAMP: 5 µg/mL, G3-YSD/LyoVec: 500 ng/mL or diABZI: 500 ng/mL) or recombinant mouse IFN-β (10 ng/mL, BioLegend, USA). Where indicated, IFNAR1 signaling was blocked using a neutralizing antibody (eBioscience™ IFNAR1 Monoclonal Antibody, clone MAR1-5A3) or respective control antibody (eBioscience™ Mouse IgG1 kappa Isotype Control, clone P3.6.2.8.1) 1 hour prior to stimulation (both: 10 µg/mL, Invitrogen, Thermo Fisher Scientific, USA). Cells were harvested at defined time points for protein analysis (3 h), gene expression (6 or 24 h) or cytokine measurements (24 h).

### Osteoblast–macrophage co-culture

A transwell-based co-culture system was used to assess indirect effects of osteoblast-derived signals on osteoclast differentiation. Osteoblasts (1 × 10^5^ cells/well) were seeded in 12-well plates overnight and pre-stimulated for 3 or 24 h. Transwell inserts (0.4 μm pore size, Sarstedt AG & Co. KG, Germany) were pre-equilibrated with medium for 30 min at 37°C. After stimulation, OB medium was exchanged to remove stimuli and transwell inserts were added. BMDMs were seeded into transwell inserts at 7 × 104 cells/insert in BMDM medium supplemented with 25 ng/mL M-CSF. RANKL (50 ng/mL) was added to induce osteoclast differentiation. Co-cultures were maintained for 4–5 days, after which BMDMs were fixed, stained and analyzed for osteoclast formation.

### TRAP staining

Osteoclast differentiation was assessed by tartrate-resistant acid phosphatase (TRAP) staining using a commercial kit (Sigma-Aldrich, Merck KGaA, Germany). Cells were fixed for 30 s at room temperature and stained for TRAP activity according to the manufacturer’s protocol (30–60 min incubation at 37°C). TRAP-positive multinucleated cells (>3 nuclei) were counted as osteoclasts and quantified by light microscopy.

### RNA isolation and quantitative PCR

Total RNA was extracted using the innuPREP RNA Mini Kit 2.0 (Analytik Jena, Germany) according to the manufacturer’s instructions. cDNA was synthesized from 500–1000 ng RNA using a reverse transcription kit (biotechrabbit GmbH, Germany). Quantitative PCR was performed using the 2x SyGreen® Mix Hi-ROX (PCR Biosystems Ltd., UK) with 400 nM primers on a StepOnePlus Real-Time PCR System (Applied Biosystems, Thermo Fisher Scientific, USA). Cycling conditions included 40 cycles with annealing/extension at 60°C. Amplification specificity was confirmed by melting curve analysis and comparison with no-reverse transcriptase (noRT) and no-template (water) controls. Gene expression levels were normalized to *Hprt1* and calculated using the 2^−ΔCq method. Primer sequences are listed in Table 1.

**Table 1.**
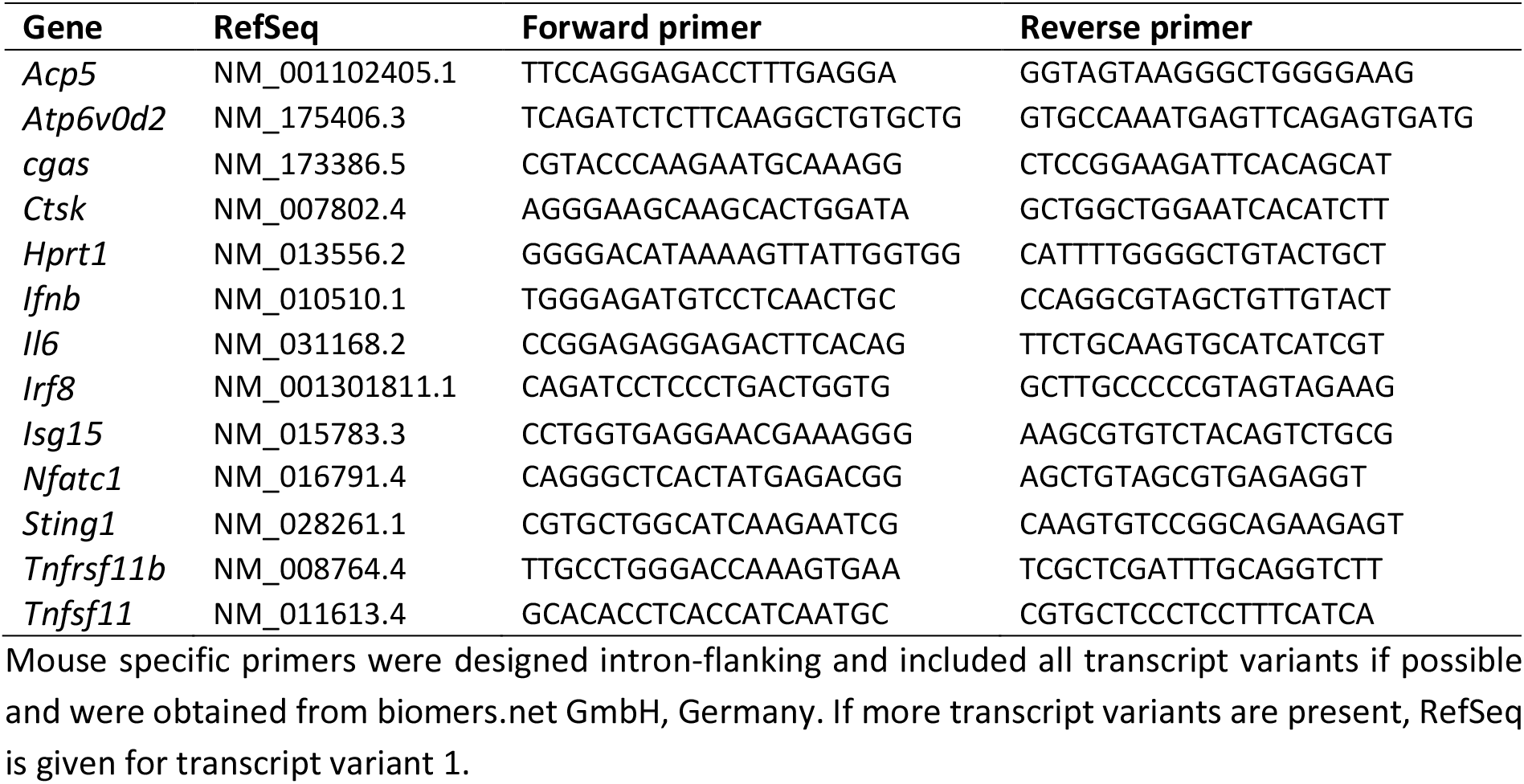
List of mouse specific oligonucleotides used for quantitative RT-PCR analysis.

For absolute quantification of cGAS transcripts, 1 × 106 RAW 264.7 cells or BMDMs were harvested. cDNA synthesis and qPCR were performed as described above. To generate a standard for absolute quantification, the cGAS amplicon was first amplified by conventional PCR and purified. A standard curve was then generated using a 10-fold serial dilution of the purified PCR product ranging from 100 ng to 1 pg. Ct values obtained from the dilution series were used to generate a linear standard curve, which was subsequently applied to calculate absolute cGAS mRNA copy numbers in unstimulated RAW 264.7 cells and BMDMs. Absolute copy numbers were determined based on the standard curve and converted to molecule numbers using Avogadro’s constant.

### Immunoblotting

Cells were lysed in RIPA buffer (50 mM Tris-HCl pH 8.0, 150 mM NaCl, 1 mM EDTA pH 8.0, 1% NP-40 (IGEPAL CA-630), 0.25% sodium deoxycholate, 1 mM Na_3_VO_4_) supplemented with protease and phosphatase inhibitors (both, SERVA Electrophoresis GmbH, Germany) for 1 h at 4 °C. Protein concentrations were determined by BCA assay (Cyanagen Srl, Italy). Protein samples were mixed with 4× SDS sample buffer (200 mM Tris-HCl pH 6.8, 8% (w/v) SDS, 40% (v/v) glycerol, 400 mM β-mercaptoethanol, bromophenol blue) and heated at 95°C for 2 min. 10 μg proteins were separated on pre-cast gradient 4–20% Tris-glycine gels (anamed Elektrophorese GmbH, Germany) at 120 V for ∼90 min and transferred to nitrocellulose membranes in transfer buffer (192 mM glycine, 25 mM Tris, 2.6 mM SDS, 0.5 mM NA_3_VO_4_, 15% (v/v) methanol) for 1 h at 2 mA/cm2. Membranes were blocked for 30 min at RT (BlueBlock PF, SERVA Electrophoresis GmbH, Germany + 0.5 mM Na_3_VO_4_), incubated with primary antibodies overnight at 4°C, followed by HRP-conjugated secondary antibodies (1 h, RT). Primary and secondary antibodies are listed in Table 2. Detection was performed using ECL (WESTAR ETA C ULTRA 2.0, Cyanagen Srl, Italy) in a ChemoStar ECL and Fluorescence Imager (Intas Science Imaging Instruments GmbH, Germany). Protein band intensities were semi-quantified from immunoblots using ImageJ software (National Institutes of Health, USA) by densitometric analysis and normalized to the respective loading control.

**Table 2.** List of antibodies used for Immunoblotting (Western Blot).

| Antibody (clone) | Source | Size (kDa) | Dilution | Company |
| --- | --- | --- | --- | --- |
| cGAS (D3O8O) | rabbit | 62 | 1:1000 | Cell Signaling Technology, USA |
| GAPDH | mouse | 36 | 1:1000 | Proteintech, USA |
| HSP90 | rabbit | 90 | 1:1000 | Cell Signaling Technology, USA |
| Phospho-IRF3 (Ser396) (4D4G) | rabbit | 45-55 | 1:1000 | Cell Signaling Technology, USA |
| NF-ATc1 (7A6) | mouse | 100-120 | 1:500 | BD Pharmingen™, BD Biosciences, USA |
| RUNX2 (D1L7F) | rabbit | 55-62 | 1:1000 | Cell Signaling Technology, USA |
| STING (D2P2F) | rabbit | 33, 35 | 1:1000 | Cell Signaling Technology, USA |
| Anti-mouse IgG, HRP-linked | horse |  | 1:1000 | Cell Signaling Technology, USA |
| Anti-rabbit IgG, HRP-linked | goat |  | 1:1000 | Cell Signaling Technology, USA |

Antibodies were all recommended for use in mouse and applied according to the manufacturer's advice. Proteins were detected by chemiluminescent luminol reaction after incubation with respective HRP-linked secondary antibody and imaged in a ChemoStar ECL Imager.

### Cytokine and secreted factor analysis

Supernatants from BMDM or LBOB gene expression experiments (24 h stimulation) were collected for analysis of secreted proteins. Cytokines released by BMDMs (IFN-β, IL-6, TNF-α and IL-10) were quantified using a LEGENDplex™ cytometric bead array (BioLegend, USA) according to the manufacturer’s instructions. Samples were analyzed at a 1:2 dilution using a MACSQuant® Analyzer 10 flow cytometer (Miltenyi Biotec, Germany). Data were analyzed using LEGENDplex™ Data Analysis Software (version 8.0). OPG (Mouse OPG ELISA Kit, Sigma-Aldrich, Merck KGaA, Germany) and IFN-β (VeriKine™ Mouse Interferon Beta ELISA Kit, PBL Assay Science, USA) were measured by ELISA according to the manufacturers’ instructions. Samples were diluted 1:20 for OPG and 1:2 for IFN-β. Standard curves were included in each assay, and all samples were analyzed in technical duplicates.

Absorbance was measured at 450 nm using a CLARIOstar® Plus microplate reader (BMG Labtech GmbH, Germany). Concentrations were calculated by fitting standard curves using a four-parameter logistic regression model.

### Flow cytometry analysis of cell surface markers and mitochondrial activity

Cell surface marker expression was analyzed by flow cytometry to assess macrophage activation and osteoblast purity. For macrophage analysis, RAW 264.7 cells and BMDMs were stained with fluorochrome-conjugated antibodies against TLR2, MHC class II, CD80 or RANK. For osteoblast characterization, cells were stained with antibodies against CD45 and CD73. All antibodies were used with 0.2 mg/ml and are listed in Table 3. Cells were incubated with antibodies for 1 h at 4°C in the dark. Mitochondrial activity of macrophages was assessed 24 h after stimulation using MitoTracker™ Green FM and MitoTracker™ Deep Red FM (100 nM, 30 min at 37°C; both Invitrogen, Thermo Fisher Scientific, USA). Data acquisition was performed using a FACSCanto™ flow cytometer (BD Biosciences, USA), and data were analyzed using FACSDiva™ software (BD Biosciences).

**Table 3.**
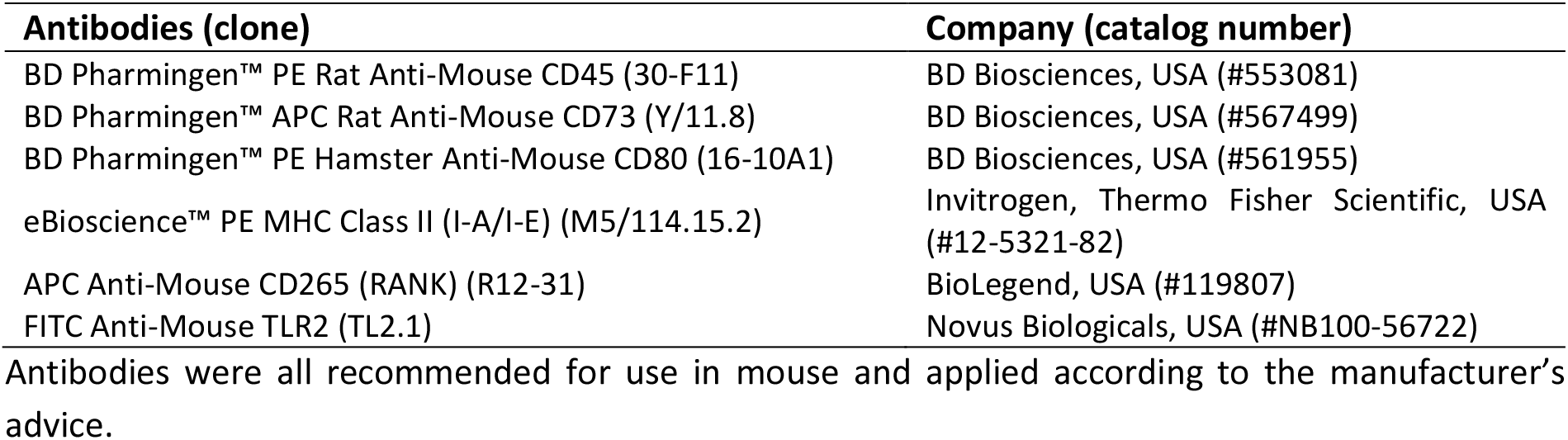
List of fluorochrome-conjugated antibodies used for flow cytometry analysis (FACS).

## QUANTIFICATION AND STATISTICAL ANALYSIS

Experiments were performed with three independent biological replicates. For RAW 264.7 cells, biological replicates represent independent experiments performed on different days using independently generated cell populations. For BMDM and LBOB experiments, each biological replicate corresponds to cells isolated from an individual mouse. Data are presented as mean ± standard error of the mean (SEM), with individual data points shown. Statistical significance was determined using ordinary one-way analysis of variance (ANOVA) followed by Bonferroni’s multiple comparisons test. A p value < 0.05 was considered statistically significant. All analyses were performed using GraphPad Prism (version 10.6.1; GraphPad Software, USA).

## ADDITIONAL RESOURCES

Not applicable.

## REFERENCES

1. Raggatt, L.J. and N.C. Partridge, Cellular and molecular mechanisms of bone remodeling. J Biol Chem, 2010. 285(33): p. 25103–8.

2. Bi, H., et al., Key Triggers of Osteoclast-Related Diseases and Available Strategies for Targeted Therapies: A Review. Front Med (Lausanne), 2017. 4: p. 234.

3. Rodan, G.A. and T.J. Martin, Therapeutic approaches to bone diseases. Science, 2000. 289 5484: p. 1508–14.

4. Khosla, S. and L.C. Hofbauer, Osteoporosis treatment: recent developments and ongoing challenges. The lancet. Diabetes & endocrinology, 2017. 5 11: p. 898–907.

5. Tsourdi, E., et al., Fracture Risk and Management of Discontinuation of Denosumab Therapy: A Systematic Review and Position Statement by ECTS. The Journal of Clinical Endocrinology & Metabolism, 2021. 106(1): p. 264–281.

6. Vargas-Franco, J.W., et al., Paradoxical side effects of bisphosphonates on the skeleton: What do we know and what can we do? Journal of Cellular Physiology, 2018. 233: p. 5696–5715.

7. Ferbebouh, M., et al., The pathophysiology of immunoporosis: innovative therapeutic targets. Inflammation Research, 2021. 70: p. 859–875.

8. Su, N., C. Villicana, and F. Yang, Immunomodulatory strategies for bone regeneration: A review from the perspective of disease types. Biomaterials, 2022. 286: p. 121604.

9. Ono, T. and T. Nakashima, Recent advances in osteoclast biology. Histochemistry and Cell Biology, 2018. 149: p. 325–341.

10. Asagiri, M. and H. Takayanagi, The molecular understanding of osteoclast differentiation. Bone, 2007. 40 2: p. 251–64.

11. Zhao, B., et al., Interferon regulatory factor-8 regulates bone metabolism by suppressing osteoclastogenesis. Nature Medicine, 2009. 15: p. 1066–1071.

12. Takayanagi, H., et al., Interplay between interferon and other cytokine systems in bone metabolism. Immunological Reviews, 2005. 208(1): p. 181–193.

13. Feng, X., RANKing Intracellular Signaling in Osteoclasts. IUBMB Life, 2005. 57(6): p. 389–395.

14. Huang, Y., et al., diABZI and poly(I:C) inhibit osteoclastic bone resorption by inducing IRF7 and IFIT3. J Bone Miner Res, 2024. 39(8): p. 1132–1146.

15. Zhang, X., X.C. Bai, and Z.J. Chen, Structures and Mechanisms in the cGAS-STING Innate Immunity Pathway. Immunity, 2020. 53(1): p. 43–53.

16. Motwani, M., S. Pesiridis, and K.A. Fitzgerald, DNA sensing by the cGAS-STING pathway in health and disease. Nat Rev Genet, 2019. 20(11): p. 657–674.

17. Ivashkiv, L.B. and L.T. Donlin, Regulation of type I interferon responses. Nature Reviews Immunology, 2013. 14: p. 36–49.

18. Wen, C., et al., Interferon signaling pathways in health and disease. Mol Biomed, 2025. 6(1): p. 135.

19. MacLauchlan, S., et al., STING-dependent interferon signatures restrict osteoclast differentiation and bone loss in mice. Proc Natl Acad Sci U S A, 2023. 120(15): p. e2210409120.

20. Lifei Liu, X.T., Mei Huang, Chenyu Zhu, Xi Chen, Yu Yuan, Samuel Bennett, Jiake Xu, Jun Zou, The effects of cGAS-Sting pathway on bone mineral density. PREPRINT (Version 1) available at Research Square 2023.

21. Gao, Z., et al., Targeting STING: From antiviral immunity to treat osteoporosis. Frontiers in Immunology, 2023. Volume 13-2022.

22. Wu, F., et al., Recent advances in targeting the cGAS-STING pathway for immunotherapy in orthopedic diseases. Front Immunol, 2025. 16: p. 1726423.

23. Boyce, B.F. and L. Xing, The RANKL/RANK/OPG pathway. Current Osteoporosis Reports, 2007. 5: p. 98–104.

24. Martin, T.J. and N.A. Sims, RANKL/OPG; Critical role in bone physiology. Reviews in Endocrine and Metabolic Disorders, 2015. 16: p. 131–139.

25. Udagawa, N., et al., Osteoclast differentiation by RANKL and OPG signaling pathways. Journal of Bone and Mineral Metabolism, 2020. 39: p. 19–26.

26. Zhang, Q., et al., Chemical regulation of the cGAS-STING pathway. Curr Opin Chem Biol, 2022. 69: p. 102170.

27. Kim, K., et al., NFATc1 induces osteoclast fusion via up-regulation of Atp6v0d2 and the dendritic cell-specific transmembrane protein (DC-STAMP). Mol Endocrinol, 2008. 22(1): p. 176–85.

28. Seebach, E., et al., Staphylococci planktonic and biofilm environments differentially affect osteoclast formation. Inflamm Res, 2023. 72(7): p. 1465–1484.

29. Takeuchi, T., et al., ISG15 regulates RANKL-induced osteoclastogenic differentiation of RAW264 cells. Biol Pharm Bull, 2015. 38(3): p. 482–6.

30. Kwon, Y., et al., Cyclic Dinucleotides Inhibit Osteoclast Differentiation Through STING-Mediated Interferon-beta Signaling. J Bone Miner Res, 2019. 34(7): p. 1366–1375.

31. Souza, P.P.C. and U.H. Lerner, Finding a Toll on the Route: The Fate of Osteoclast Progenitors After Toll-Like Receptor Activation. Front Immunol, 2019. 10: p. 1663.

32. Kim, J., et al., Lipoproteins are an important bacterial component responsible for bone destruction through the induction of osteoclast differentiation and activation. Journal of Bone and Mineral Research, 2013. 28(11): p. 2381–2391.

33. Yu, Z.C., et al., The STING inhibitor C-176 attenuates osteoclast-related osteolytic diseases by inhibiting osteoclast differentiation. FASEB J, 2023. 37(4): p. e22867.

34. Huang, L., et al., Osteoclastogenesis Responds to STING Inhibition in a Non-Monotonic Manner. FASEB J, 2026. 40(6): p. e71596.

35. Weinstock-Guttman, B., et al., Interferon-beta modulates bone-associated cytokines and osteoclast precursor activity in multiple sclerosis patients. Mult Scler, 2006. 12(5): p. 541–50.

36. Tsukasaki, M., et al., OPG Production Matters Where It Happened. Cell Rep, 2020. 32(10): p. 108124.

37. Cawley, K.M., et al., Local Production of Osteoprotegerin by Osteoblasts Suppresses Bone Resorption. Cell Rep, 2020. 32(10): p. 108052.

38. Choe, C.H., et al., Transmembrane protein 173 inhibits RANKL-induced osteoclast differentiation. FEBS Lett, 2015. 589(7): p. 836–41.

