## Supplementary material for "cGAS–STING induced IFN-β acts as a dual regulator of osteoclastogenesis via direct and osteoblast-mediated mechanisms": Suppl. Material

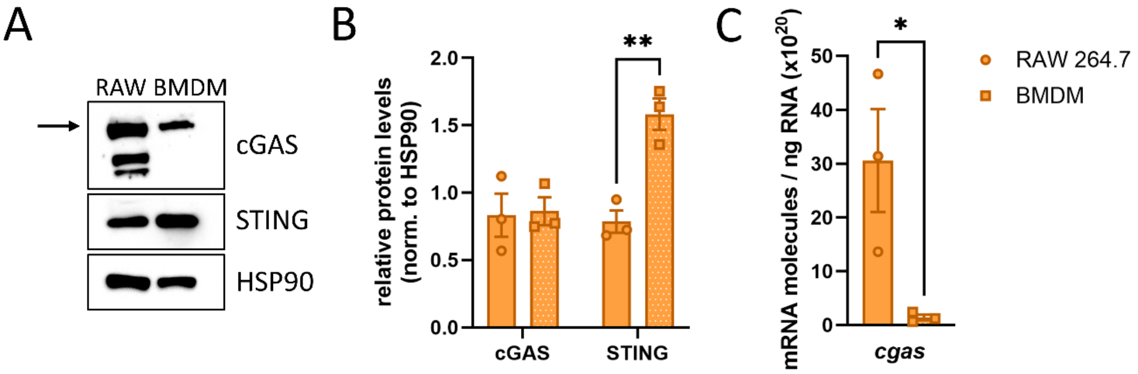


Suppl. Figure S1. Expression levels of cGAS–STING in macrophages. (A) Immunoblot analysis of cGAS and STING protein levels in BMDMs and RAW 264.7 cells. Equal amounts of total protein (10 µg per sample) were loaded for each condition. HSP90 served as a loading control, although its expression differed between the two macrophage cell types. (B) Quantification of cGAS and STING protein levels in BMDMs and RAW 264.7 cells. (C) Absolute qPCR quantification of cGAS mRNA copy numbers in BMDMs and RAW 264.7 cells. (A-C) BMDMs were cultured in the presence of 25 ng/mL recombinant mouse M-CSF throughout all experiments. Bar graphs show mean ± SEM with individual data points. Statistical analysis was performed using one-way ANOVA with Bonferroni post hoc test (n = 3).

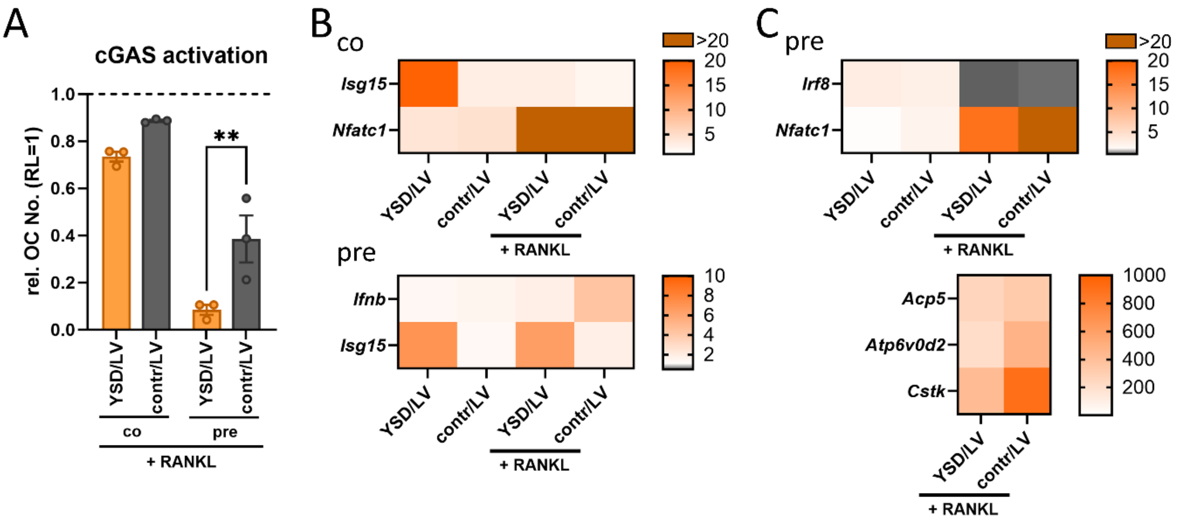


Suppl. Figure S2. cGAS pre-stimulation affects osteoclastogenesis in RAW 264.7 macrophages. Effect of cGAS activation by G3-YSD in comparison to YSD control, both complexed in LyoVec™ (500 ng/mL) on RANKL-mediated osteoclast formation of RAW 264.7 cells. (A) Quantification of relative osteoclast numbers per well in RAW 264.7 cells after co-stimulation of G3-YSD / control and RANKL (co) or pre-stimulation with G3-YSD / control for 24 hours and subsequent RANKL (pre). Osteoclast numbers per well are shown relatively to the RANKL control. (B+C) Gene expression analysis of interferon-related genes and osteoclast-associated genes 48 h after stimulation with

50 ng/mL RANKL. (B) shows *Isg15* and *Nfatc1* expression levels of co-stimulation (top) and *Ifnb* and *Isg15* expression levels of pre-stimulation (bottom). (C) shows osteoclast-associated genes of pre-stimulation. (A-C) Data are normalized to the unstimulated control. Heatmaps display mean values. RL: RANKL; LV: LyoVec™ transfection agent; contr: YSD control.

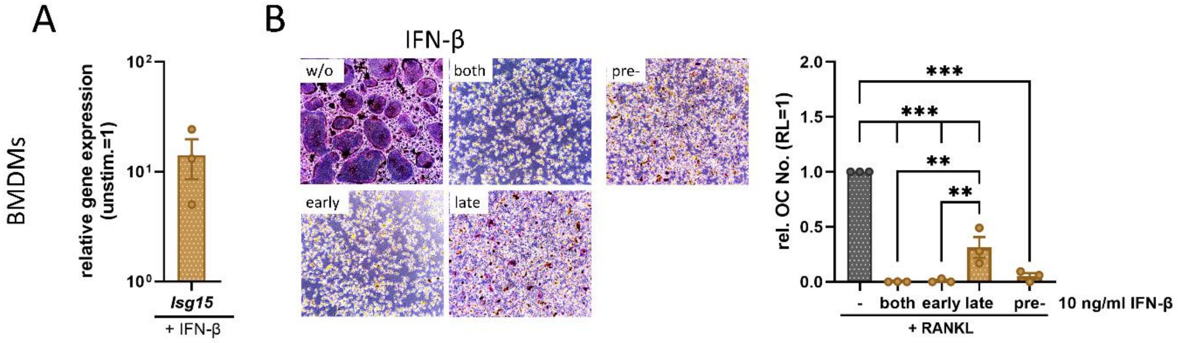


<u>Suppl. Figure S3. Direct effect of IFN-β on osteoclastogenesis.</u> (A) Gene expression analysis of IFN-responsive gene *Isg15* in BMDMs after stimulation with IFN-β (recombinant mouse protein, 10 ng/mL) for 3 h. (B) Effect of IFN-β (recombinant mouse protein, 10 ng/mL) on RANKL-mediated osteoclast formation of BMDMs. Cells were treated with IFN-β throughout differentiation ("both"), during early stages (first 3 days) or during late stages (days 3–5/6). For pre-treatment IFN-β was added 24 h prior to RANKL stimulation. In this case, the cytokine was removed before 50 ng/mL RANKL was added. Representative images of osteoclasts (left) and quantification of relative osteoclast numbers per well (right). (A+B) BMDMs were cultured in the presence of 25 ng/mL recombinant mouse M-CSF throughout all experiments. Osteoclast numbers per well are shown relatively to the RANKL control. Bar graphs show mean ± SEM with individual data points. Statistical analysis was performed using one-way ANOVA with Bonferroni post hoc test (n = 3).

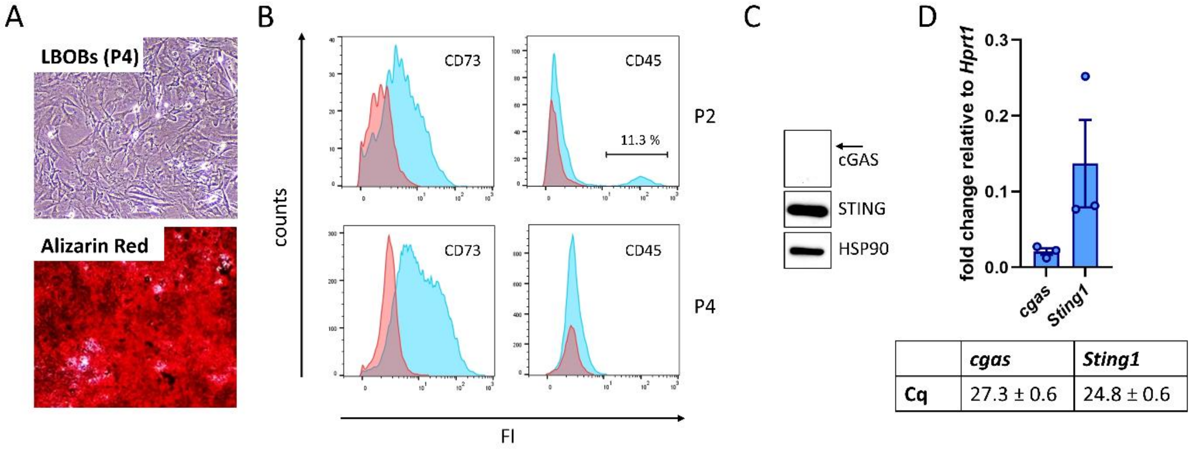


| | *cgas* | *Sting1* |
|---|---|---|
| **Cq** | 27.3 ± 0.6 | 24.8 ± 0.6 |

<u>Suppl. Figure S4. Characterization of osteoblasts isolated from long bone fragments (LBOBs).</u> (A) shows cell morphology and Alizarin Red staining to assess mineralization. (B) Surface marker expression of CD73 and CD45. (C) Immunoblot analysis of cGAS and STING protein levels. HSP90 served as a loading control. (D) RT-qPCR data of *cgas* and *Sting1* expression levels normalized to reference gene *Hprt1*. Cq values are listed in the table below. Bar graph shows mean ± SEM with individual data points. (A-D) Data are representative for n=3 independent LBOB populations.

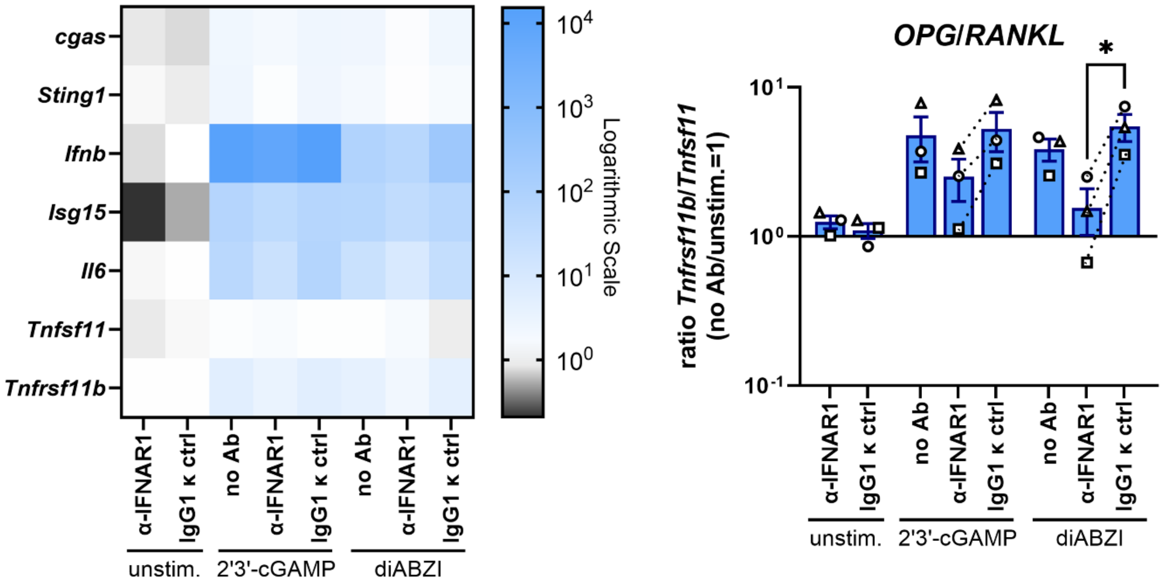


Suppl. Figure S5. Inhibition of IFNAR signaling affects STING-mediated changes of OPG–RANKL axis in osteoblasts. Gene expression analysis of osteoblasts after neutralization of IFNAR1 (10 µg/mL) and stimulation with STING agonists (2’3’-cGAMP: 5 µg/mL, diABZI: 500 ng/mL) for 6 h. Left: heatmap of cGAS–STING pathway-associated genes, including *Tnfsf11* (RANKL) and *Tnfrsf11b* (OPG). Right: ratio of *Tnfrsf11b* to *Tnfsf11* mRNA levels. Data are normalized to the unstimulated, no-antibody (no Ab) control. Heatmaps display mean values, and bar graphs show mean ± SEM with individual data point symbols indicating independent experiments, respectively. Statistical analysis was performed using one-way ANOVA with Bonferroni post hoc test (n = 3). Ctrl: control; Ab: antibody.
